# EEG markers of drowsiness do not predict lapses in attention during sleep deprivation

**DOI:** 10.1101/2023.10.02.560530

**Authors:** Sophia Snipes, Elias Meier, Simone Accascina, Reto Huber

## Abstract

During drowsiness, maintaining consistent attention becomes difficult, leading to behavioral lapses. Oscillation bursts in the electroencephalogram (EEG) might predict such lapses, since alpha bursts increase with inattention and theta bursts increase with time spent awake. However, while lapses increase with time awake, paradoxically alpha bursts decrease, and the behavioral relevance of theta bursts is unknown. Therefore, we investigated whether theta or alpha bursts predicted lapses either when well rested (baseline, BL) or sleep deprived (SD). EEG was measured in 18 young adults performing the lateralized attention task, and the timing of bursts was related to trial outcomes (fast, slow, and lapse trials). Against expectations, neither theta nor alpha bursts were more likely during lapses, either at BL or SD. Both were more likely before fast trials, but only at BL. Fast and slow trials were followed by decreases in both theta and alpha bursts, but the effect was reduced during SD, partially due to local increases in bursts. Considering prior literature, these results indicate that bursts have a non-linear relationship with vigilance. The lack of a direct relationship to lapses may be because bursts originate from task-unrelated areas, and the differences during SD suggest these areas change with time awake.

## INTRODUCTION

Sleepiness can be deadly. Just 17 h of extended wake has the same effect as mild alcohol intoxication and up to 20% of road traffic accidents are attributed to insufficient sleep (Dawson & Reid, 1997; Gibbings et al., 2022; Horne & Reyner, 1995). While multiple cognitive systems are likely compromised during sleep deprivation, the most affected seems to be sustained attention (Lo et al., 2012). Laboratory tests of sustained attention such as the psychomotor vigilance task (PVT) reliably capture increasing behavioral lapses with time spent awake, circadian rhythm, and even cumulative sleep restriction (Basner & Dinges, 2011; Dinges & Powell, 1985; Graw et al., 2004; Van Dongen et al., 2003). Given the role sleepiness can have on health and safety, there is a justifiable interest in identifying the neural mechanisms leading to these behavioral lapses.

One of the most notable features of brain activity are bursts of oscillations in the EEG, reflecting different states of vigilance (Schomer & Silva, 2011). Since the first EEG recordings, theta oscillations (4-8 Hz) have been associated with sleepiness (Smith, 1938), and alpha oscillations (8-14 Hz) with inattention and “neuronal idling” (Adrian & Matthews, 1934; Klimesch et al., 1998). While theta activity increases with time spent awake (and therefore drowsiness), curiously alpha activity decreases (Aeschbach et al., 1997; Cajochen et al., 2002; Finelli et al., 2000; Snipes et al., 2023; Strijkstra et al., 2003). Nevertheless, within a recording session, alpha activity has been found to predict behavioral lapses and therefore track fluctuations in vigilance (Huang et al., 2007; Makeig & Jung, 1996).

Such oscillations have typically been quantified using spectral power (Aeschbach et al., 1997; Cajochen et al., 2002; Finelli et al., 2000; Klimesch et al., 1998; Snipes et al., 2022) or in the case of theta as single waves (Andrillon et al., 2021; Bernardi et al., 2015; Fattinger et al., 2017; Hung et al., 2013). However, recently we were able to automatically quantify bursts directly (Snipes et al., 2023). In resting EEG, with increasing time awake we found a near-linear increase in the quantity of theta bursts and decreases in alpha bursts (which matches the literature), but independent increases in oscillation amplitudes for both theta and alpha. Given this dissociation, we aimed to re-examine the relationship between EEG bursts and behavioral outcome and determine whether the presence of either a theta or an alpha burst was predictive of a lapse in a visual attention task. Furthermore, we wished to determine whether the increase in theta bursts with time spent awake could explain the increase in attentional lapses and whether the relationship between bursts and outcome differed during sleep deprivation compared to baseline.

We collected high-density EEG data from 18 young healthy adults undergoing a 4/24 extended wake paradigm (4 h of sleep, 24 h of wake; full schedule in Figure 1A). To capture attentional lapses, we used an adaptation of the PVT, the Lateralized Attention Task (LAT). Like the PVT, this involved fixating on a rectangle in the center of the screen, with stimuli appearing every 2-10 seconds. However, the stimuli were faint circles that would shrink within 0.5 s (Figure 1B). Participants had to push a button whenever they saw the stimulus, and it would flash green if caught in time. A lapse was defined as any trial for which no response was given. To further increase the proportion of lapses, the task was performed under soporific conditions with lights off and seated in a comfortable armchair with foot and headrest. The LAT was performed three times when well-rested (baseline, BL), and three times following at least 20 h of extended wake (sleep deprivation, SD).

**Figure 1:**
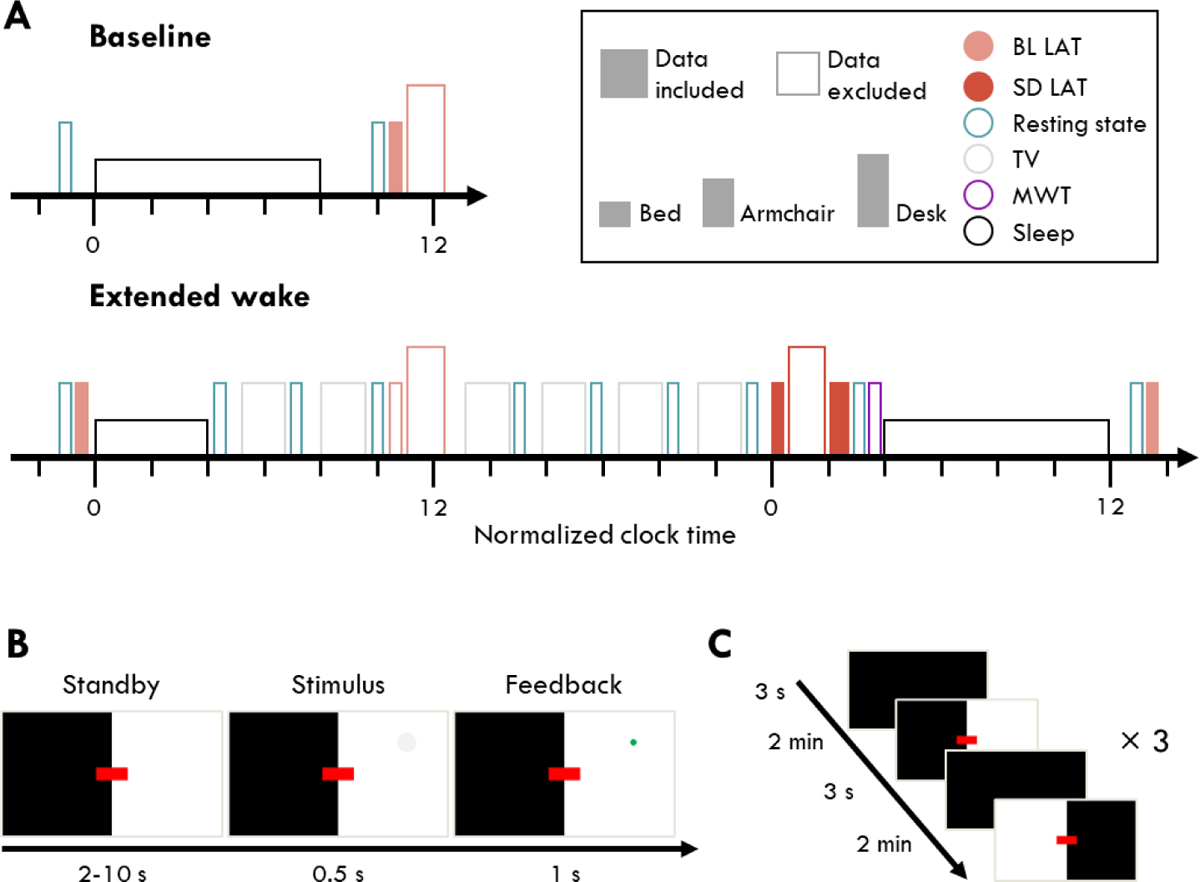
Study design. A: Experiment schedule. Each block indicates an EEG recording session. Filled blocks indicate data analyzed in this paper. Color indicates the activity participants engaged in: gray, watching TV; turquoise, the resting state recordings (analyzed in Snipes et al. (2023)); peach, baseline (BL) task blocks; red, sleep deprivation (SD) task blocks; purple, the maintenance of wakefulness test (MWT); black, sleep. The height of each block indicates the experimental condition in which data was collected: short, lying in bed; medium, seated in a comfortable armchair with foot and backrest; tall, seated at a desk, analyzed in Snipes et al. (2022). Brief empty spaces indicate transition periods allowing for delays. Six longer breaks were included prior to each TV block in which participants were provided with meals. Circadian time was normalized across participants to their habitual bedtime. Participants at baseline and during the recovery night were free to wake up when they wished, and at the beginning of the extended wake period they were woken up after 4 h of sleep. B: Lateralized Attention Task (LAT) trial. Participants had to fixate on the red rectangle, and every 2-10 seconds, a gray circle would appear somewhere in the white area. If they pressed a button before the stimulus completely shrunk away, it would flash green as positive feedback. C: The 12-minute LAT consisted of 6 blocks, alternating between the left and right screen being illuminated. Brief pauses separated the switch.

To test the hypothesis of whether bursts could contribute to lapses, we investigated both the occurrences of bursts in time (proportion of channels with a burst at every timepoint, i.e. globality) and the occurrences in space (the proportion of timepoints with a burst in a given interval for each EEG channel) around stimulus onset. We expected that bursts could be associated to lapses in two ways: they could be more likely exactly when the stimulus was present, indicating that bursts disrupted processing and responding to the stimulus; or they could be uniformly more likely in the seconds around lapse trials, indicating a general marker for a non-vigilant state (Makeig & Jung, 1996) even if the bursts themselves do not directly cause the lapse. By analyzing the changes in burst occurrences with high temporal resolution, we could determine the direction of causality. By analyzing changes in topography, we had greater sensitivity to local effects.

To provide a comparison with a reliable lapse-causing event, we also applied the same analysis to eyeclosures, which during sleep deprivation reflect a mixture of blinks, resting with eyes closed, and genuine brief episodes of sleep known as microsleeps (Hertig-Godeschalk et al., 2020; Ong et al., 2013). Furthermore, we repeated the analyses controlling for trials with eye-closures, in order to exclude any lapses (and bursts) associated with such obvious causes.

## RESULTS

### The LAT is sensitive to attention lapses

We first determined whether the LAT was an appropriate task for measuring lapses following extended wake, and whether the adaptations from the PVT effectively increased eyes-open lapses. Participants performed one BL PVT session the morning after the baseline night of sleep, and one SD PVT session after 20 h of wake, counterbalanced with the first LAT. The PVT defines lapses as trials with reaction times (RT) > 0.5 s. This means that lapses include both 1) sluggish responses and 2) “true” lapses in attention when the participant misses the stimulus onset, then recovers later eventually noticing the counter, and presses the button (Figure 2A). Capturing both is what makes the PVT a sensitive and robust measure of sleepiness; however this is suboptimal for investigating whether a given event can cause an attention lapse. Shifting the lapse threshold to higher RT values avoids these sluggish responses, but this also increases the proportion of lapses with eyes closed (Figure 2C). Therefore, for lower RTs the PVT cannot distinguish between slow responses and lapses, and for higher RTs lapses mostly reflect microsleeps.

**Figure 2:**
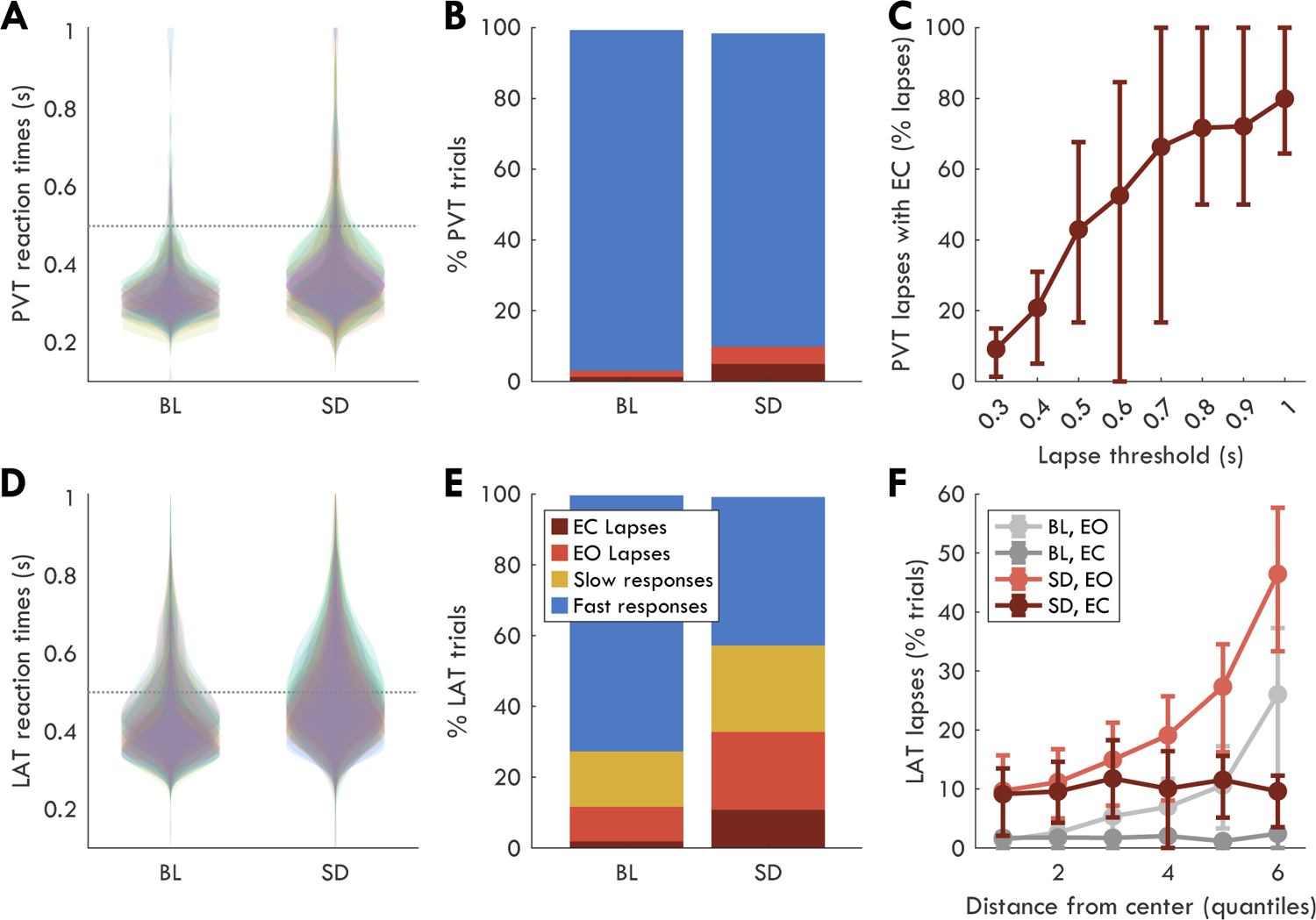
LAT versus PVT behavioral outcome measures. A: Reaction times during the PVT. Each colored “violin” represents the distribution of an individual participant (N=18). The dotted horizontal line at 0.5 s marks the threshold over which the trial was considered a lapse. B: Average distribution of PVT trials based on response outcome (fast responses: RT < 0.5 s; lapses: RT > 0.5 s) with lapses split by whether eyes were open or closed (N=13; due to eye tracking data loss. Only includes participants with data in both session blocks). EC fast trials are not included in the bar graph (they are the sliver of whitespace at the top). C: Percentage of PVT lapses that are with EC, depending on the RT threshold used to define lapses (N=9). Error bars indicate interquartile range around the average. The higher the RT threshold, the more lapses are due to EC. Anderson et al. (2010) performed a similar analysis, although they found 10% of lapses were with EC at 0.5 s cutoff, which only increased to 90% after ∼2 s. This difference may be due to our soporific conditions. D: Same as A for LAT (N=18), although the LAT was performed 3 times for each session block instead of just once. N.B: BL recordings are pooled from three different days, whereas SD recordings were within the same 3-4 h timespan, and therefore also include time-on-task effects (Doran et al., 2001). E: Same as B for LAT (N=17). The LAT distinguishes slow trials as 0.5 s < RT < 1 s (yellow) and lapses as trials where no response was given. F: Percentage of LAT trials that are lapses, split into 6 quantiles based on the distance from the fixation point, such that 1 is closest and 6 is furthest (N=17). 100% indicates all trials in that quantile were lapses (either EO or EC). Gray indicates BL trials, red SD trials. Darker shades reflect EC, lighter shades EO. Error bars indicate the interquartile range.

The main difference between the LAT and PVT is that LAT stimuli appeared only briefly and are difficult to detect, minimizing the orienting response (Pavlov, 1927). This means that even a brief lapse in attention results in missing the stimulus entirely. Therefore, with the LAT it is possible to separate slow responses from complete lapses in attention (Figure 2D-E). In this experiment, the average percentage of LAT trials classified as a lapse was 12% [interquartile range: 5, 19] at BL, and 33% [22, 44] at SD. By comparison, PVT lapses were 3% [1, 4] of BL trials, and 10% [2, 14] of SD trials (Figure 2B). Only 4% [1, 6] of PVT SD trials were eyes-open lapses, and 22% [13, 30] of LAT trials. This was a highly significant increase from BL, when only 10% [5, 16] of LAT trials were eyes-open lapses (N = 17, t = 6.31, p < .001, g = 1.47). Therefore, the LAT captured substantially more eyes-open lapses in attention than the PVT, making it the more appropriate task to evaluate the potential detrimental effects of a given neuronal event.

Because stimuli appeared at any distance from the fixation point during the LAT, it could be that most lapses occurred at the edge of participants’ visual field. To check if this was the case, trials were divided into 6 quantiles based on radial distance from the fixation point (Figure 2F). At BL, considering only EO trials, the closest quantile had 1.3% [0.0, 1.8] of trials as lapses and the furthest had 26.0% [10.4, 37.3]. Therefore, while distance was clearly a major contributor of lapses at BL, there was no distance after which stimuli were completely missed and were thus outside of the field of view. In other words, distance from fixation increased the chances of a lapse but didn’t determine one.

### Bursts capture the majority of periodic power

EEG oscillatory bursts were detected with cycle-by-cycle analysis (Cole & Voytek, 2019). Replicating what we found during resting EEG in the same participants (Snipes et al., 2023), extended wake increased the total number of theta bursts (Figure 3A) and decreased alpha bursts (Figure 3C) across all channels. At BL theta bursts were on average 0.80 s [0.75, 0.83] long, with amplitudes of 15 μV [13, 16], and during SD they were 0.86 s [0.78, 0.88] long, and 18 μV [15, 19]. Alpha bursts were 0.57 s [0.50, 0.60] at BL and 0.56 s [0.49, 0.60] during SD, with amplitudes of 13 μV [11, 14] and 14 μV [12, 15] respectively.

**Figure 3:**
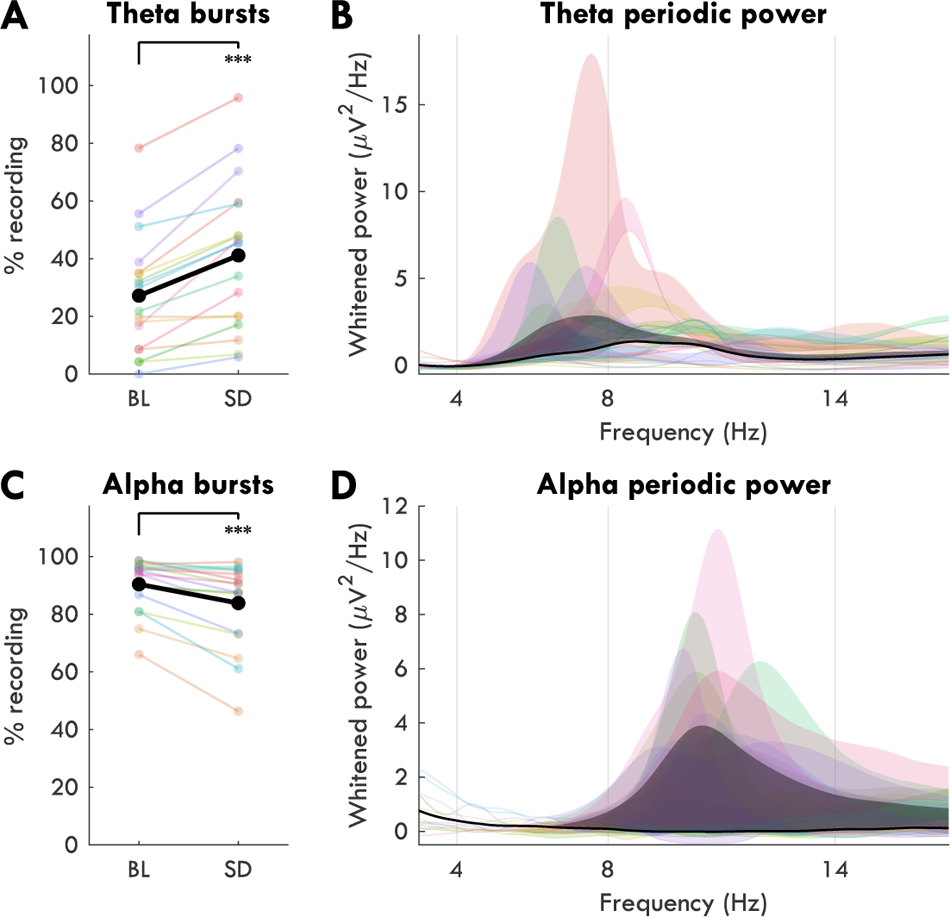
Burst detection. A: Percentage of the task recording characterized by theta at baseline (BL) and after sleep deprivation (SD). Thin colored lines indicate individual participants, black lines indicate the group average. Paired t-tests were conducted with α = 5%, such that: * p < .05, ** p<.01, *** p < .001. B: Periodic component of the power spectra from frontal channels during SD, before and after timepoints containing theta bursts were removed from the signal. For each colored patch, the bottom line indicates power without theta bursts, and the patch reflects power with theta bursts. The black patch indicates the average across participants. Spectra were “whitened” by subtracting the 1/f “colored” component (see methods). Channels included were the front region of interest described in Snipes et al. (2022). C-D: Same as A-B, but for alpha during BL, from posterior channels.

To evaluate the successfulness of the burst detection, we compared whitened EEG power spectra before and after removing theta bursts from frontal channels during SD (Figure 3B), and removing alpha bursts from posterior channels during BL (Figure 3D). Whitened spectra were used to focus on the changes in the periodic component of the EEG (driven by oscillations), and avoid the aperiodic 1/f “background” activity (Donoghue et al., 2020). On average, removing theta bursts reduced theta power by 57% [40, 79] and removing alpha bursts reduced alpha power by 100% [N = 16, IQR: 98, 102].

### Bursts predict faster trials at baseline, but show no relationship to trial outcome during sleep deprivation

To determine whether there was a temporal relationship between bursts and lapses, we calculated the globality of bursts across time with a window from 2 seconds before to 4 seconds after stimulus onset, split by trial outcome (fast, slow, lapses). Values were z-scored to the mean and standard deviation of burst globality across the recordings of that session block (BL, SD) to minimize interindividual variability. We conducted paired t-tests for each timepoint relative to the average burst globality present in the session block (0 after z-scoring), with false-discovery rate (FDR) correction for multiple comparisons.

To validate this analysis, we first applied it to the relationship between eye-closures and trial outcomes during BL (Figure 4A) and SD (Figure 5A). During BL, the only timepoints during lapse trials that had p < .05 were during the stimulus window, peaking 0.16 s (max time) from stimulus onset (N = 13, t = 4.53, pfdr = .002, g = 1.70). This clearly reflects blinks occurring exactly when the stimulus was shown. By contrast, eye-closures during the stimulus window of fast trials were substantially less likely than average (max t: 0.30 s; N = 18, t = −14.59, pfdr < .001, g = −4.72), as well as in the seconds before fast trials (max time: −0.44 s; N = 18, t = −16.06, pfdr < .001, g = −5.19). For slow trials, eye-closures were also significantly less likely before the stimulus, albeit with smaller effect sizes (max time: −1.36 s; N = 16, t = −6.91, pfdr < .001, g = −2.36) and during the stimulus presentation (max time: 0.28 s; N = 16, t = −8.71, pfdr < .001, g = −2.98). Notably, there was an increase in the likelihood of eye-closures exactly at stimulus onset relative to fast trials, indicating that blinks exactly when the stimulus appeared contributed to the delayed response.

**Figure 4:**
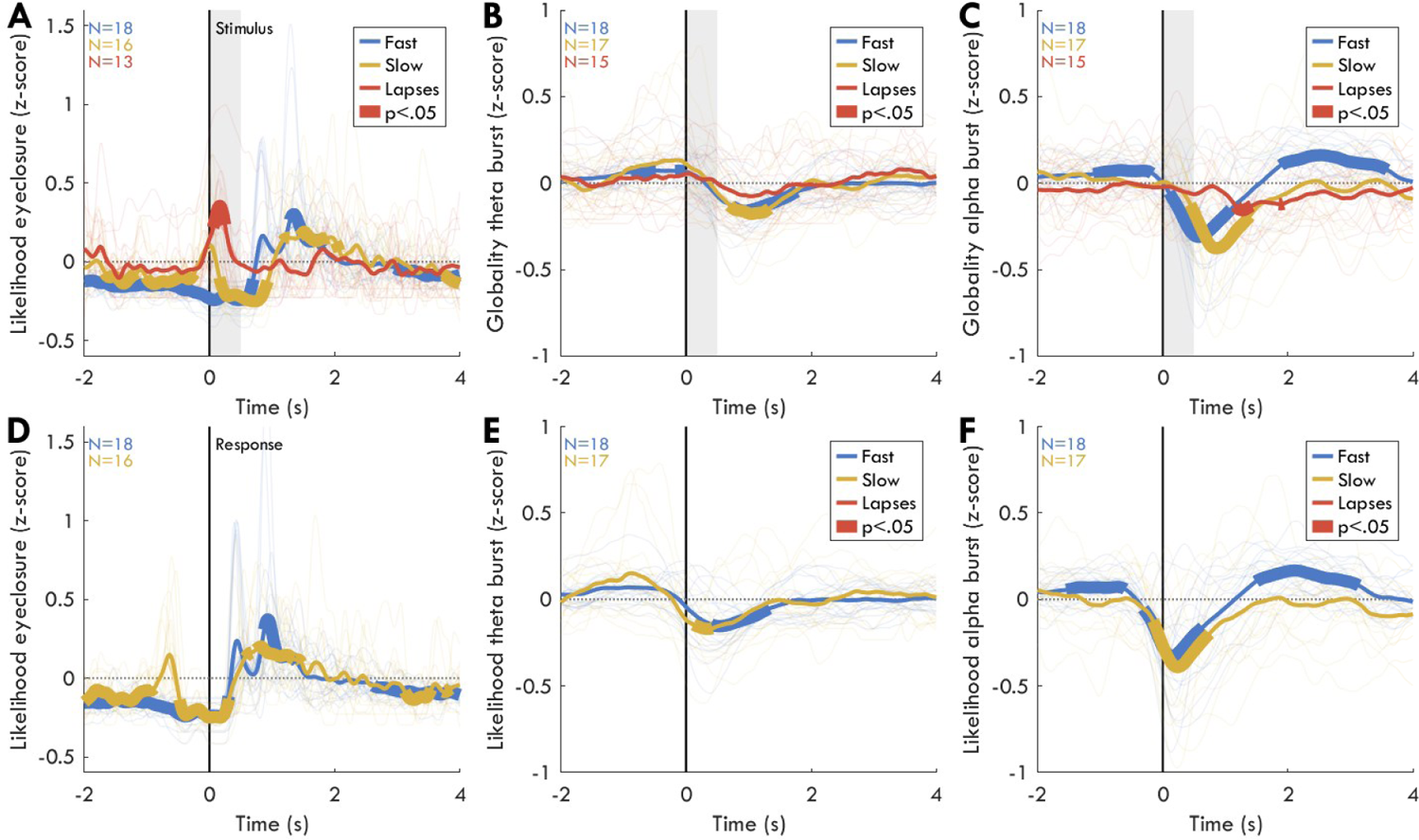
Distribution in time of bursts and eyes-closed relative to trial outcomes during baseline recordings. A: Difference in proportion of trials with EC for each outcome type relative to the recording average amount of EC, such that values at 0 (dotted horizontal line) represents no difference from the average. The thick vertical line represents stimulus onset, and the gray patch the time in which the stimulus was visible. Light thin colored lines represent individual averages, medium lines indicate the group average, and thick segments reflect timepoints in which the difference was statistically significant, with p< .05, FDR corrected for multiple comparisons. Sample sizes are indicated in the top left. The average amount of time spent with eyes closed was 6% [4, 8] for the session block. B: Same as A, but for theta bursts. Trials during which eyes were closed during the stimulus window were excluded. The average globality was 2% [0, 2]. C: Same as B for alpha bursts. The average globality was 17% [11, 20]. D-F: Same as A-C, with trials locked to the response instead of stimulus onset. N.B., the fast and late negative peaks are more aligned in F than in C, indicating that the desynchronization is more related to the response than to stimulus processing.

**Figure 5:**
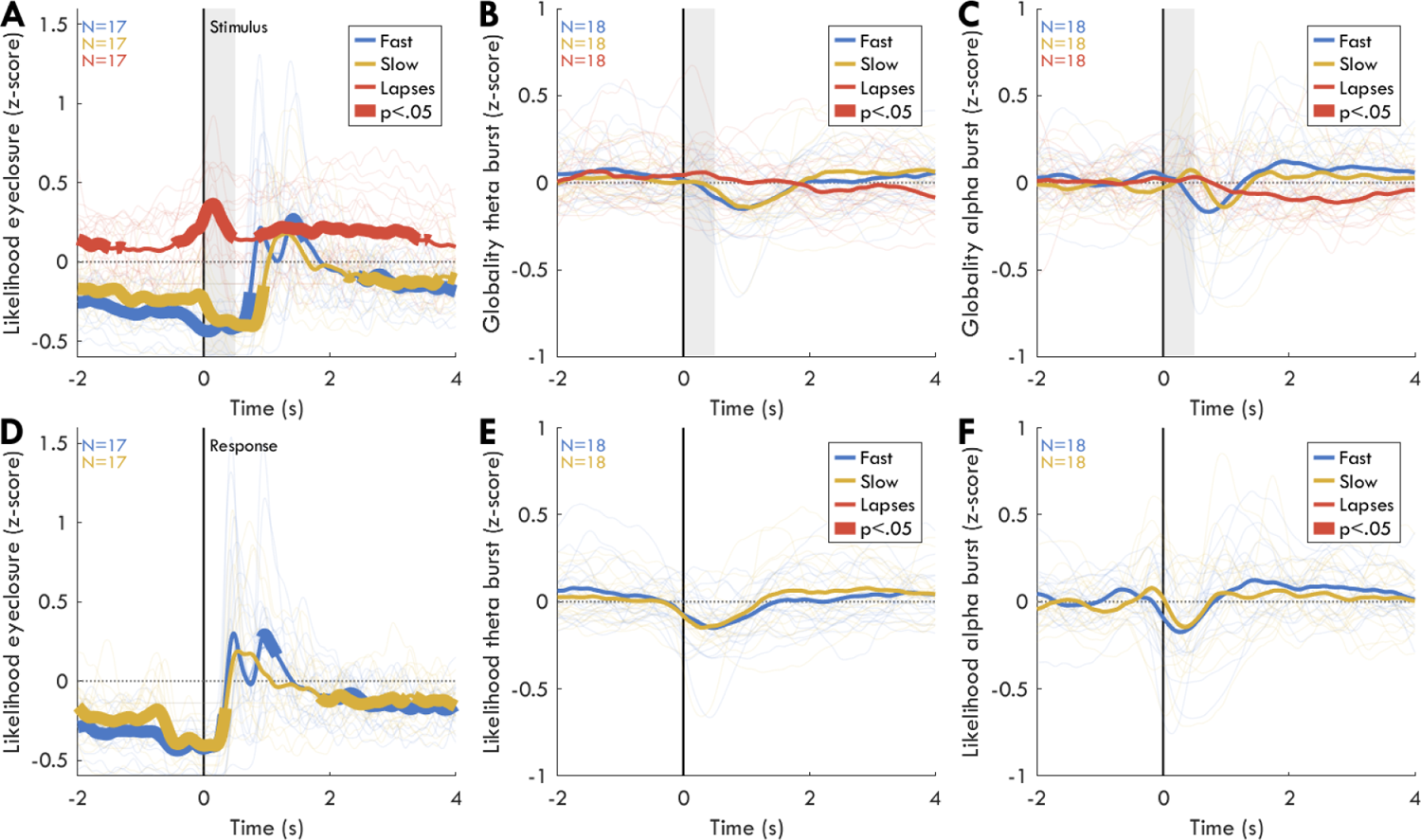
Distribution in time of eyes-closed and bursts relative to trial outcomes during sleep deprivation. Same as Figure 4. A,D: eyes closed. The average time spent with eyes closed was 17% [14, 21] of the session block. B,E: theta bursts. The average globality was 5% [1, 5]. C,F: alpha bursts. The average globality was 12% [7, 16]. N.B. for lapses and slow trials, the N is lower (indicated in top left) because some participants did not have enough of such trials to be included in this analysis. The N is different between A and B,C because the burst analyses excluded trials with EC.

During SD (Figure 5A), the only major difference from BL was that eye-closures were more likely in the seconds prior to lapses (max time: −1.89 s; N = 17, t = 3.20, pfdr = .011, g = 1.06) and after lapses (max time: 1.90 s; N = 17, t = 3.68, pfdr = .005, g = 1.22). This likely reflects the fact that, in addition to blinks during the stimulus, long stretches of eye-closures also occurred and resulted in lapses. Altogether, this analysis clearly reflects how eye-closures are both a direct cause of lapses (peaking during the stimulus window), as well as an indicator of an overall non-responsive state (likely reflecting microsleeps). Thus, similar dynamics could be expected of EEG bursts.

At BL for lapse trials (Figure 4B), there was no timepoint for which theta burst globality differed from average either before (max time: −2.00 s; N = 15, t = 0.86, pfdr = .713, g = 0.30), during (max time: 0.04 s; N = 15, t = 1.11, pfdr = .606, g = 0.39) or after the stimulus appeared (max time: 3.55 s; N = 15, t = 1.84, pfdr = .320, g = 0.65). Instead, theta bursts were significantly more likely before the stimulus appeared for fast trials (max time: −0.09 s; N = 18, t = 3.26, pfdr = .049, g = 1.06), followed by decreases after the stimulus window (max time: 1.47 s; N = 18, t = −9.55, pfdr < .001, g = −3.09), with the nadir after the response (Figure 4E), before returning to average levels. For slow trials, the pattern was the same albeit not significant before trials (max time: −0.74 s; N = 17, t = 2.47, pfdr = .137, g = 0.82), but after (max time: 1.03 s; N = 17, t = −3.99, pfdr = .021, g = −1.33). Altogether, this indicates that increases in theta burst globality before trials was predictive of improved performance. It’s peculiar how late trials have on average higher theta globality prior to stimulus onset than fast trials, even if the effect is not significant. This seems driven by 3 participants with elevated theta at stimulus onset, and may indicate that the relationship between bursts and behavior is not consistent across individuals.

For alpha bursts at BL (Figure 4C), the pattern was similar albeit more pronounced. There were no significant differences in alpha burst globality during lapse trials before (max time: −1.06 s; N = 15, t = −1.50, pfdr = .315, g = −0.53) or during the stimulus window (max time: 0.30 s; N = 15, t = −0.76, pfdr = .636, g = −0.27), however the likelihood of alpha bursts gradually decreased across the trial window, becoming significant > 1 s after stimulus onset (max time: 1.27 s; N = 15, t = −3.18, pfdr = .028, g = −1.12). Like for theta, fast trials were anticipated by more alpha bursts (max time: −0.90 s; N = 18, t = 3.62, pfdr = .013, g = 1.17), which rapidly decreased across the stimulus window, peaking just after the response (max time: 0.40 s; N = 18, t = −4.92, pfdr = .003, g = −1.59). This corresponds to the well-known event-related desynchronization of alpha (Schomer & Silva, 2011), and likely explains the same observed for theta. There was an additional positive rebound in alpha bursts after the trial (max time: 2.40 s; N = 18, t = 5.95, pfdr = .001, g = 1.92). Slow trials did not have significantly more alpha bursts before the stimulus (max time: −1.44 s; N = 17, t = 1.55, pfdr = .294, g = 0.52), but had the same desynchronization (max time: 0.97 s; N = 17, t = −5.10, pfdr = .003, g = −1.69), albeit slightly delayed.

During SD, no timepoint for any trial outcome survived correction for multiple comparisons for either theta bursts (Figure 5B) or alpha bursts (Figure 5C). Theta bursts and lapses remained non-significant before (max time: −1.52 s; N = 18, t = 1.83, pfdr = .475, g = 0.59), during (max time: 0.30 s; N = 18, t = 1.30, pfdr = .603, g = 0.42) and after the trial (max time: 4.00 s; N = 18, t = −2.67, pfdr = .181, g = −0.86). Theta wasn’t even trending higher before fast trials (max time: −1.29 s; N = 18, t = 1.83, pfdr = .475, g = 0.59), although the response desynchronization was trending (max time: 1.01 s; N = 18, t = −3.60, pfdr = .065, g = −1.16), as it was for slow trials (max time: 1.14 s; N = 18, t = −4.11, pfdr = .065, g = −1.33). The gradual decrease in alpha bursts during lapse trials remained, albeit not significant (max time: 2.76 s; N = 18, t = −3.45, pfdr = .149, g = −1.12). The higher alpha globality before fast trials was also no longer significant during SD (max time: −0.17 s; N = 18, t = 2.48, pfdr = .149, g = 0.80), and neither was the desynchronization (max time: 0.78 s; N = 18, t = −2.86, pfdr = .149, g = −0.93).

This analysis was further repeated excluding timepoints with eyes closed, to control for lapses clearly driven by blinks or microsleeps. For theta bursts at BL (Suppl. Figure 4-1B), the increased theta globality before the stimulus was only trending (max time: −0.06 s; N = 18, t = 3.18, pfdr = .089, g = 1.03), as was the desynchronization for slow trials (max time: 0.89 s; N = 16, t = −3.19, pfdr = .089, g = −1.09), otherwise the results were comparable. For alpha bursts at BL (Suppl. Figure 4-1C), the only difference was that alpha was significantly less likely for lapse trials also before the stimulus (max time: −0.99 s; N = 12, t = −3.18, pfdr = .034, g = −1.24). During SD (Suppl. Figure 5-1B), no timepoint was even trending for either theta or alpha bursts (pfdr >= .118).

### Widespread desynchronizations after fast trials at baseline are reduced or even locally reversed during sleep deprivation

The previous analysis was conducted by pooling bursts from all channels and may have masked local effects. We therefore conducted the same analysis for bursts detected in each channel, collapsing time into four windows: Pre (−2 to 0 s from stimulus onset), Stimulus (0 to 0.3 s from stimulus onset), Response (0.3 to 1 s from stimulus onset), and Post (2 to 4 s from stimulus onset). Paired t-tests for each channel compared the proportion of theta or alpha bursts split by trial outcome for the different time windows, relative to the averages of each channel. FDR correction was applied for each topography to correct for multiple comparisons across channels.

During BL, for theta bursts (Figure 6A) no channel during any time window for any trial outcome was statistically significant after FDR correction. However, matching Figure 4B, there was more theta prior to fast trials across the entire topography, and less theta for fast trials in the response window. Instead for alpha bursts (Figure 6B), there was significantly more alpha before the stimulus of fast trials in midline occipital-parietal channels (significant in 17% of channels; max channel: 71, N = 18, t = 5.24, pfdr = .004, g = 0.13). During the stimulus window, 50% of channels had significantly less alpha, peaking bilaterally over lateral occipital-parietal areas (max ch: 96; N = 18, t = −6.62, pfdr < .001, g = −0.37). Given the timecourse in Figure 4C, this very likely reflects the stimulus-evoked desynchronization, which begins almost immediately after stimulus onset. This desynchronization spreads to 78% of channels during the response window, peaking over left parietal areas (max ch: 59; N = 18, t = −7.53 pfdr < .001, g = −0.80). This was followed by a widespread positive rebound over 74% of channels (max ch: 67; N = 18, t = 5.54, pfdr = .002, g = 0.21). Slow trials only had the response desynchronization in 56% of channels (max ch: 60; N = 17, t = −9.52, pfdr < .001, g = −0.84).

**Figure 6:**
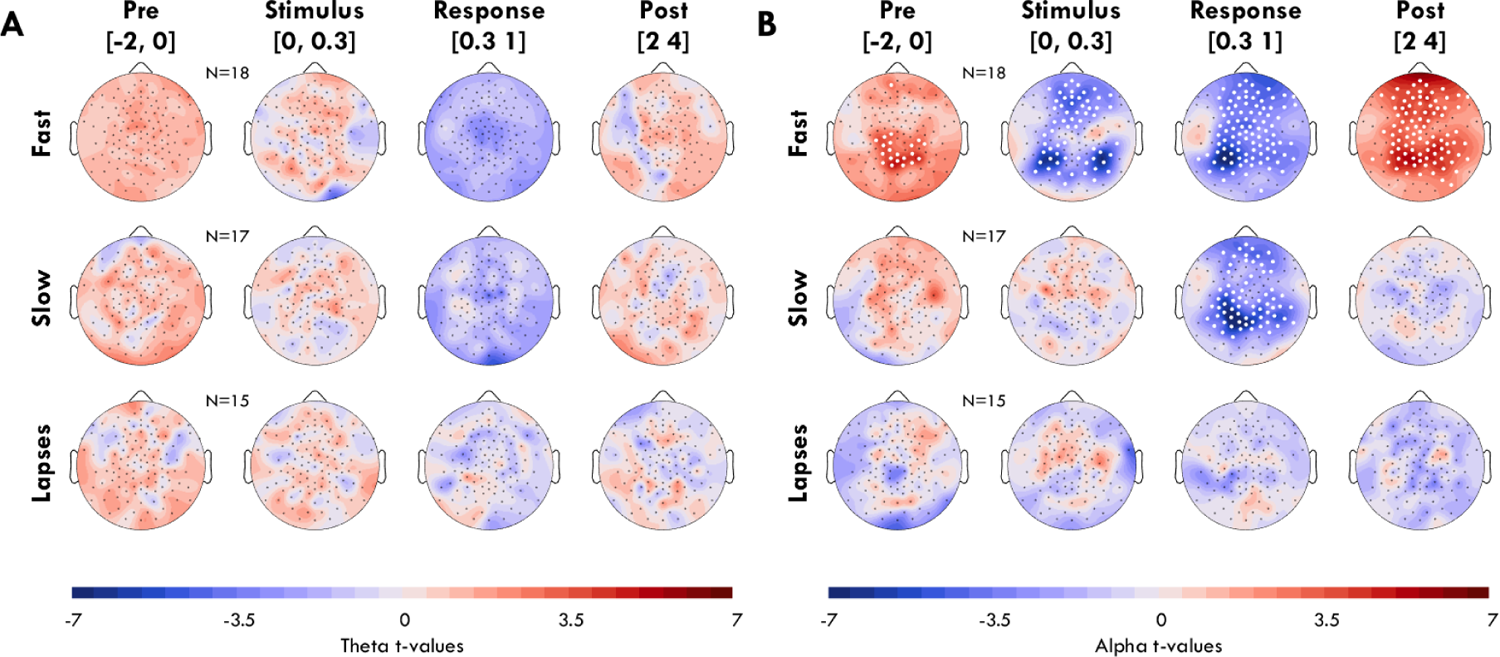
Topography of burst proportions per trial type during baseline. A: Difference in theta burst proportion from session average for different trial outcomes (rows) and time windows (columns), with the seconds ranges indicated in square brackets. Color indicates t-values comparing the proportion of theta for a given outcome in a given time window to the recording average for a given channel, such that red indicates more theta relative to the recording average. White dots indicate statistically significant differences, p < .05. FDR correction was applied for each topography. Sample size is indicated for each trial outcome. B: Same for alpha.

During SD, again no channel showed significant changes in theta bursts (Figure 7A). Instead, alpha bursts (Figure 7B) decreased significantly in the Post window of lapse trials in 25% of channels (max ch: 62; N = 18, t = −6.35, pfdr < .001, g = −0.25). This is extremely peculiar and could possibly reflect an increase in attention after too much time has passed between stimuli, or the presence of lapses just at the onset of sleep, in which case the decrease in alpha during post reflects the presence of a microsleep. The fact that this effect was not present at BL could simply be a lack of power, as there were fewer lapses and therefore fewer participants (BL N = 15, SD N = 18). For fast trials, the pre-stimulus and stimulus windows no longer had any significant channels, and the response window had reduced decreases in alpha, occupying only 27% of channels (max ch: 66; N = 18, t = −8.44, pfdr < .001, g = −0.62), and notably also significant increases in alpha in a single left temporal channel. The rebound after the trial was also reduced, and significant in 26% of channels (max ch: 96; N = 18, t = 4.71, pfdr = .011, g = 0.32). Repeating the analysis with only eyes-open trials and timepoints did not substantially change the results (Suppl. Figure 6-1, 7-1), although 2 channels were still significant before fast trials during SD.

**Figure 7:**
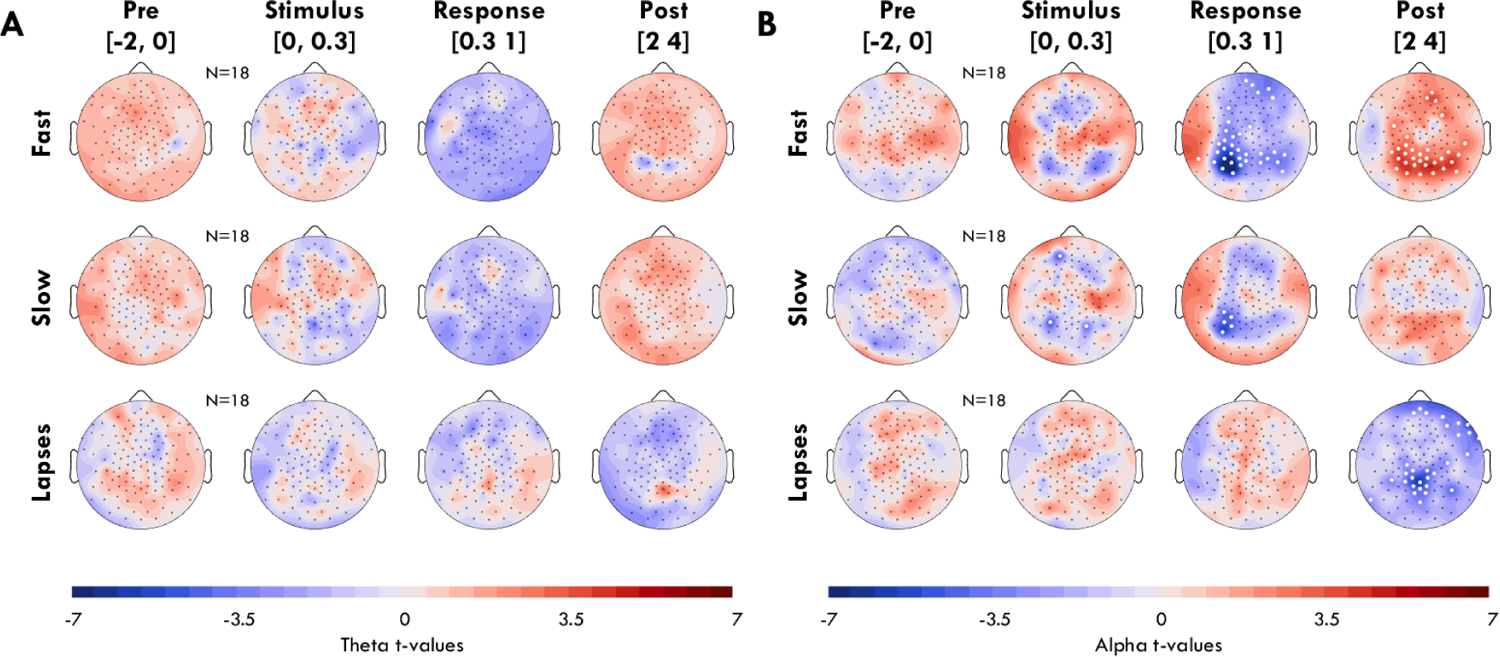
Difference in burst proportion by trial window and outcome during sleep deprivation. Same as Figure 6. A: Theta bursts. B: Alpha bursts.

To better understand what could drive the reduced effect at SD, we inspected the individual difference topographies of the Response window of fast trials for alpha bursts, as this had the largest overall effects (Figure 8, Suppl. Figure 8-1, 8-2). It was first of all notable how at BL, while most participants displayed widespread decreases in alpha, a couple showed equal or greater increases, especially outside the parietal areas (P08, P13). It’s possible this reflects an earlier rebound than for the others. However, during SD, more participants showed increases in alpha outside of parietal areas (P05), whereas others merely reduced the intensity of the desynchronization (P01, P16). Therefore, the change in topography during the response window of fast trials was due to both a reduction in desynchronization, and an increase in alpha bursts in other cortical areas.

**Figure 8:**
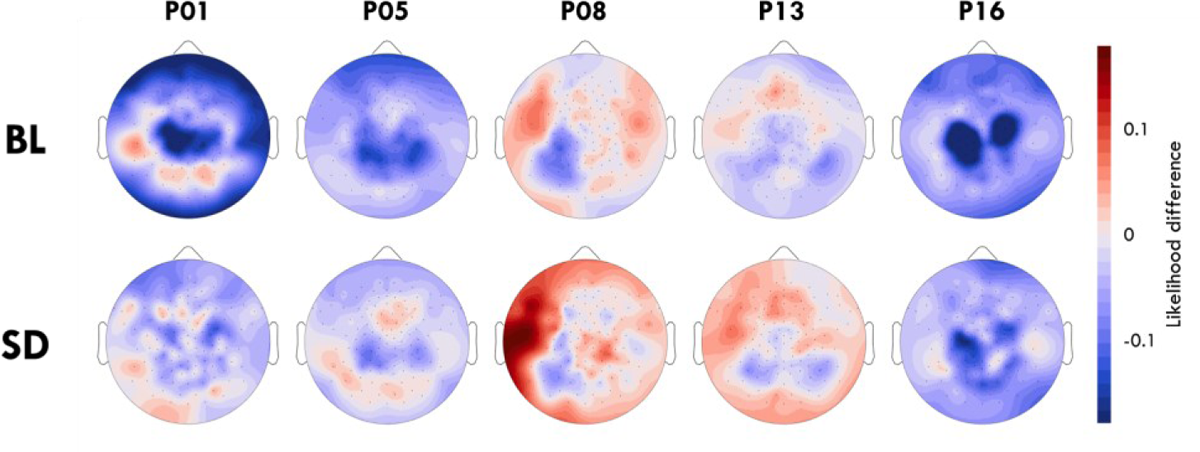
Individual participants’ change in alpha in the Response window of fast trials. Color indicates the difference between the proportion of timepoints containing alpha (averaged across trials) and the average proportion of alpha across the entire session block for that participant, such that red indicates above average alpha for that channel. The plots are scaled individually. See Suppl. Figure 8-1 and 8-2 for all participants.

## DISCUSSION

With this study, we wanted to determine whether there was any meaningful relationship between oscillation bursts (considered markers of reduced vigilance) and attentional lapses. We also wanted to determine whether the relationship between bursts and trial outcome differed when sleep deprived compared to well rested. Our results yielded three main findings. First, we found no relationship between either theta or alpha bursts and lapses that could indicate a causal relationship. Neither were more likely to occur *during* the stimulus window of lapse trials (or even slow trials) so bursts couldn’t be a direct cause. Also, neither theta or alpha bursts occurred *before* lapse trials, so bursts weren’t predicting within-session reductions in vigilance. This was the case both at baseline and following extended wake. Second, we actually found the opposite at baseline: elevated theta and alpha bursts prior to fast trials. This would imply instead that bursts reflect periods of higher vigilance, at least when well rested, contradicting the literature (Hanslmayr et al., 2011; Makeig & Jung, 1996; Sauseng et al., 2005). Third, we found that the burst dynamics of fast trials were severely diminished during sleep deprivation, due in part to the appearance of opposite dynamics in select regions. This indicates that the relationship between bursts and behavior can differ substantially depending on time spent awake.

### How can alpha anticipate better performance?

The elevated amount of alpha bursts prior to fast compared to slow and lapse trials (Figure 4C) directly contradicts previously published results identifying alpha power as a within-session marker of reduced vigilance (Hanslmayr et al., 2011; Makeig & Jung, 1996; Sauseng et al., 2005). For example, in a study by Huang et al. (2007), well-rested participants performed a sustained attention driving task with similar “game mechanics” to the PVT and LAT (occasional driving adjustment required every 3 to 7 s). When sorting trials into low and high driving performance, the authors found that low performance “drowsy” trials were anticipated by higher alpha power, followed by a negative and then positive deflection, neither of which was present for alert trials. We found the complete opposite, with the faster trials at baseline showing higher pre-stimulus alpha globality, and stronger post-stimulus fluctuations. If anything, alpha bursts were even less likely during lapse trials than average.

An explanation for these opposing results is that alpha and vigilance could follow a non-linear relationship, such that low alpha characterizes both extremely high and extremely low vigilance. Recent work by Pfeffer et al. (2022) found in fact that alpha activity and pupil diameter follow this inverted U pattern, such that both small pupils (indicating low vigilance) and large pupils (indicating high vigilance) are associated with lower alpha compared to intermediate values. It could be that previous studies like those of Huang et al. were conducted under higher overall vigilance, from which drops in alertness (and performance) corresponded to increased alpha. Instead in our study, participants were already quite drowsy at baseline: the LAT had slower reaction times than the PVT (Figure 2B), participants indicated higher subjective sleepiness ratings during this task (Snipes et al. [2022]), and they performed the LAT under deliberately soporific conditions (dark room, armchair). This would explain why for this study, further drops in alertness even at baseline resulted in *decreased* alpha. Therefore, the pre-stimulus alpha across these two studies follows the same inverted U relationship described in Pfeffer et al. and can explain our seemingly contradictory results. The fact that alpha has a non-linear relationship with vigilance also makes sense given that alpha decreases with sleep deprivation, so it cannot be a monotonic marker of alertness.

### What do theta bursts do?

We do not know what theta bursts are for, but an important step towards arriving at an answer is determining whether they are qualitatively distinct from alpha bursts or not. In support of the idea that they are different is the fact that they originate from nearly opposite ends of the brain, preferentially occur during different mental activities, and change in opposite directions with increasing time awake (Klimesch, 1999; Schomer & Silva, 2011; Snipes et al., 2023). Therefore, while alpha may follow an inverted-U relationship to vigilance, theta rather increases linearly with increasing time awake and therefore sleepiness. However, since we did not find theta to predict either behavioral lapses or late responses, even during sleep deprivation, theta may not reflect the loss of vigilance with time awake, but some yet-unknown process.

In support of theta and alpha having the same function, we found they had similar dynamics around fast trials (Figure 4). Even more compelling, the occurrence of both theta and alpha are associated with reduced fMRI BOLD (functional magnetic resonance imaging, blood oxygen level dependent) activity from the regions generating the oscillations (Scheeringa et al., 2008, 2012), and these sources weren’t involved in the ongoing task (Kirschfeld, 2005; Michels et al., 2010; Rihs et al., 2007; Scheeringa et al., 2009; Snipes et al., 2022). Therefore, they could both be a form of idling rhythm for disengaged cortical areas. Since every brain region has a preferred resting state frequency (Frauscher et al., 2018), it may be that the *only* difference between theta and alpha is the source of the oscillation, which alters their frequency but not their function. If both oscillations reflect disengagement in task-unrelated areas, this would explain why they have no direct impact on behavior, but may reflect similar shifts in vigilance following task stimuli. However, this does not explain why alpha would decrease with time spent awake while theta increases.

### Why do burst dynamics change with sleep deprivation?

There is a possible explanation for how theta and alpha could have the same idling function while still demonstrating independent changes with time awake: sleep deprivation results in a rearrangement of which brain areas are being used for the same mental activity. This was discovered with fMRI studies, which found that the same task during sleep deprivation involved partially different cortical networks than at baseline (Chee & Choo, 2004; Drummond et al., 2005). It is therefore possible that the changes in burst dynamics likewise reflect a rearrangement in which areas are preferentially idling vs involved in the task. In general, it could be that alpha-generating areas are more often idling at baseline, whereas theta-generating frontal areas idle more during sleep deprivation. What it actually means for a brain region to be “idling” is of course an open question; here we use the term because of the association between oscillations and BOLD reductions, but these bursts could still be functionally relevant (Klimesch, 2012; Schomer & Silva, 2011). Regardless, a rearrangement in which regions tend to oscillate could explain many of the differences between baseline and sleep deprivation, for example the increase in left-temporal alpha following behavioral responses during sleep deprivation (Figure 7B, Figure 8).

Rihs et al. (2009) found that the modulation of alpha during tasks was dependent on the baseline levels of alpha activity for that participant, with higher baselines predicting greater modulation. This could apply to individual brain regions as well. Therefore, at baseline, a subset of areas is more likely to be involved in the task, another subset more likely to be idling, and the presence of a stimulus and response causes the latter areas to desynchronize and then rebound. Instead during sleep deprivation, the areas that used to produce the most alpha now idle less, and therefore desynchronize less following stimulation. At the same time, areas that may have been more engaged at baseline now idle more, but these different areas may desynchronize and synchronize at different speeds, thus appearing to follow opposite dynamics. This explanation involves a lot of additional assumptions, but our finding that the topography of EEG bursts changes during sleep deprivation is compatible with those from fMRI showing the use of different task networks. This hypothesis could therefore be tested with simultaneous EEG and fMRI during sleep deprivation.

### What could cause behavioral lapses?

Bursts could still cause behavioral lapses; it may just be that our methods were not sufficiently sensitive. There are probably more brain areas *not* involved in such a simple task than those that are. Therefore, even if bursts were equally likely everywhere, it’s possible that the perfect timing of a burst in a task-relevant area when a stimulus is present is too rare to be noticeable with this analysis. It may even be so localized that it is not apparent when looking at bursts from the scalp. Then in practice, bursts are *not* equally likely everywhere, and it may be that the regions producing the most prominent bursts are not the ones most related to this task.

Another popular hypothesis is that local sleep may be behind such lapses (Bernardi et al., 2015; Hudson et al., 2020). Local sleep in the form of isolated slow waves intruding on wake was also found to produce changes in theta power (Vyazovskiy et al., 2011). It’s still possible that this local sleep theta is related to behavioral lapses, but given how the majority of theta power was captured in bursts (Figure 3), it wouldn’t make sense to consider all changes in theta power with sleep deprivation to be a manifestation of local sleep. This still means that the most obvious events in the EEG aren’t necessarily “doing” something that we can observe, and vice versa, whatever actually causes critical attentional lapses remains undetected.

### Limitations

An important limitation of this study is sample size. With 18 participants, we only have enough power for medium-large effect sizes. Lapse trials in particular were sometimes too few per condition, so this reduced the sample size even further. There also seems to be large interindividual differences both in topography and the timing of bursts related to trial outcomes, and therefore subpopulations may yield different results. It may be necessary to focus on more localized brain areas, which could require looking at intracortical data in patients. Alternatively, a longer time awake may result in sufficiently intense/numerous theta bursts that a relationship with behavior could emerge. Therefore, we cannot definitively conclude that bursts do not predict lapses at all, just that if the effect is there, it is not particularly strong.

It is also important to note that the method we used for the burst detection differs from prior studies that relied on spectral power; it discounts differences in amplitude, although amplitude and globality are somewhat correlated. Therefore, differences from previous results are to be expected and require careful interpretation. In our previous analysis of bursts in resting EEG, we found a topographical dissociation between amplitudes and quantities of alpha bursts, such that amplitudes were higher in lateral occipital areas (matching typical alpha power topographies), and quantities higher in more central-parietal areas (Snipes et al., 2023). This means the results in this manuscript could be more sensitive to the more numerous, but smaller parietal alpha rhythms than studies using power. It could also be that amplitudes are a better indicator of vigilance than the mere presence/absence of a burst. All the same, the alpha dynamics we observe are remarkably similar to those reported from time-frequency power analyses (Huang et al., 2007; Nir et al., 2017), even if associated with different trial outcomes.

## Conclusion

The main conclusion of this paper is that EEG oscillatory bursts are not major contributors to behavioral lapses, nor do they always predict within-session reductions in vigilance, either when well rested or sleep deprived. The reason bursts don’t directly affect performance could be that both alpha and theta bursts tend to originate from task-unrelated areas and therefore don’t usually conflict with behavior. The reason bursts don’t consistently reflect either reduced or increased vigilance could be because the major sources of such oscillations change with the overall level of vigilance, most notably following sleep deprivation. Ultimately, if there is a neural marker anticipating behavioral lapses which is detectable in the scalp EEG, it is likely substantially more subtle than the large oscillations readily observed in the raw signal.

## METHODS

Different data from this experiment has previously been reported in Snipes et al. (2022) where the overall study design, participant selection, and EEG preprocessing was established, and in Snipes et al. (2023) where the burst detection method was developed and reported. These previous publications were exploratory analyses conducted in order to refine the analysis pipeline and better understand the increase in theta power observed during sleep deprivation. The data analyzed in this paper was deliberately set aside for this manuscript to test the hypothesis of whether a given EEG signal could explain behavioral lapses.

### Participants

18 participants completed the experiment. University student applicants were screened for good health, good sleep quality, and at least some sleep deprivation vulnerability. 19 participants were recruited, and one participant dropped out midway. Mean age was 23 ± 1 years old, 3 were left-handed, all had normal or corrected-to-normal vision, and self-reported no hearing impairments. Data collection and interaction with participants was conducted according to Swiss law (Ordinance on Human Research with the Exception of Clinical Trials) and the principles of the Declaration of Helsinki, with Zurich cantonal ethics approval BASEC-Nr. 2019-01193.

### Experiment design

The full experimental schedule is depicted in Figure 1A. Participants came to the laboratory for two experimental bouts: baseline, and extended wake. During the baseline, participants went to bed at their habitual bedtime, and were free to wake up whenever they chose. On average they slept 8.0 ± 0.5 h. In the morning, they were provided breakfast and had at least 40 minutes from when they woke up to when they began task recordings. During the extended wake bout, participants slept only 4 hours, were kept awake 24 h, alternating between watching TV, rest recordings (Snipes et al., 2023), and breaks. We refer to this as a *4/24 extended wake* paradigm.

The main experiment task block consisted of 6 counterbalanced tasks performed at a computer desk at three timepoints (tallest blocks in Figure 1A): the morning after the baseline night, the same time during extended wake, and after 20 h of extended wake (Snipes et al., 2022). These task blocks included the LAT and PVT. However, these two tasks were also performed under soporific conditions: seated in an armchair with footrest and headrest, lights turned off, with the task projected onto a wall. These soporific recordings are the ones analyzed in this manuscript. The soporific LAT and PVT were performed in counterbalanced order with the desk task blocks, and then additionally the evening before and the morning after the extended wake bout. The soporific LAT was then performed two more times after the last task recording of SD. PVT data reported in Figure 2 comes from the one baseline and one sleep deprivation soporific recordings. The LAT BL session block was composed of the recordings from baseline, evening before, and morning after the extended wake bout, marked in pink in Figure 1A. N.B. these were at three different circadian times: mid morning, evening, and midday. The LAT SD session block was composed of the first counterbalanced recording after 20 h of wake, and the final two repetitions, marked in red in Figure 1A. Therefore, for half the participants, the three SD LAT tasks were performed after more than 22 h awake, back to back, whereas for the other half, the first SD LAT (and SD PVT) was performed around 20 h awake, before the 2 h computer task block.

### The Lateralized Attention Task

The LAT is a 12 min visual-spatial reaction time task, modeled after the PVT (Basner & Dinges, 2011). 6 blocks (2 min each) alternated between having the left or right visual hemifield in white, and the other in black (Figure 1B). Participants had to maintain fixation on a red rectangle in the center of the screen, and covertly attend to the white half of the screen. Every 2-10 s a feint grey circle (1 cm radius, #F7F7F7) would appear randomly in any location of the illuminated hemifield and shrink away completely within 0.5 s. While faint, the stimuli were still above detection threshold levels for all participants when presented near the fixation point. Participants were instructed to press a button (on a MilliKey button box) before the circle disappeared, in which case the circle would freeze and flash green. Responses earlier than 0.1 s were considered false alarms. Responses from 0.5 to 1 s after the circle completely disappeared were considered late. If 5 stimuli were missed consecutively, an alarm would sound to wake up the participant (this occurred at least once for almost every participant during SD). During the delay periods, 50 ms pink noise tones were presented every 1.5-5 s at ∼50dB. Participants were instructed to ignore these tones. Overall, participants had between 100-130 trials per recording, and therefore usually >300 trials per session block. One participant completed only one SD LAT recording, another participant completed only two SD LAT recordings. All others had 3 BL and 3 SD EEG recordings. For each analysis at every timepoint, there had to be at least 15 trials involved per trial outcome.

The 0.1 s and 1 s cutoffs for considering trial responses was decided a-priori, based on typical adult reaction times in the PVT, in order to exclude false-alarms. While there may have been a few RTs slower than 1 s, during sleep deprivation 99% of RTs were within 830 ms. Compared with the ∼33% of trials resulting in a lapse, this clearly indicates that the vast majority of lapses would have happened regardless of the 1 s cutoff.

### EEG analysis

EEG data was recorded at 1000 Hz with Cz reference, using BrainAmp amplifiers and 128 channel EGI Geodesic EEG nets. Preprocessing and data analysis were done using EEGLAB (Delorme & Makeig, 2004) and custom MATLAB scripts (R2019b, R2022b, R2023a). Data was downsampled to 250 Hz and broadband filtered between 0.5-40 Hz. Major artifacts were identified visually, and physiological arti-facts (eye movements, heartbeat, muscle activity) were removed with independent component analysis (ICA). The final channel count was 123. Further details are provided in Snipes et al. (2022).

Bursts were detected using cycle-by-cycle analysis (Cole & Voytek, 2019), with adaptations previously published (Snipes et al., 2023). To identify bursts, first clean EEG data was filtered in narrow overlapping bands 4 Hz wide, from 2 to 16 Hz, and for each band, zero-crossings were identified. Between zero-crossings, positive and negative peaks were identified in the broadband filtered data, and a cycle was defined from positive to positive peak. Each cycle was characterized by properties (e.g. period, amplitude consistency with neighboring peaks), and a minimum number of consecutive cycles had to have properties meeting a set of criteria to be classified as a burst. Multiple criteria sets were used, on both the signal and its inverse, and so from bursts overlapping in time in the same channel, only the longest was kept. Bursts that were only present in one channel were also excluded.

Bursts were then sorted as theta and alpha based on the mean negative peak-to-peak period. In our previous publication, cycles had to meet a single set of criteria to be considered a burst. Here, we chose three sets of criteria which together captured a larger fraction of oscillatory activity. These were identified through trial-and-error on independent data (PVT and resting EEG), with the goal of capturing as much of the oscillatory signal that could be visually identified.

One criteria set had many criteria with low thresholds. At least 4 consecutive cycles had to have a period in the narrowband filtered range, similar consecutive periods (period consistency = .5), similar consecutive amplitudes (amplitude consistency = .4), similar rising and descending amplitudes (flank consistency = .5), they had to correlate with neighboring cycles (shape consistency = .2; centering the negative peaks, neighboring cycles were correlated, and the smallest correlation was considered), a minimum proportion of *timepoints* that changed amplitude in the correct direction of the cycle (monotonicity in time = .6), a minimum proportion of the *amplitude* that changed in the correct direction of the cycle (monotonicity in amplitude = .6) and a maximum ratio of the largest segment in the incorrect direction (reversal ratio = .6; ratio of the largest reversal to the smallest cycle flank).

The second criteria set aimed to detect short but “smooth” bursts. At least 3 consecutive cycles had to have a period in the narrow band range, a period consistency of .7, a flank consistency of .3, and monotonicity in amplitude of .9.

The third criteria also had fewer criteria than the first, but stricter requirements. There was no requirement for the period to be within the narrowband range, but 5 consecutive cycles had to have period consistency, amplitude consistency, monotonicity in amplitude and flank consistency all above .6.

The success of the burst detection was evaluated by comparing the power spectrums of EEG data with data after timepoints containing bursts had been removed (Figure 3B,D). Power was calculated using Welch’s method, with 8 s windows and 50% overlap. Participants were excluded if there was less than 1 minute of data without bursts. Given that the EEG spectra measure both periodic (oscillations) and aperiodic activity (1/f “colored background noise”), we subtracted the aperiodic component from the spectra, thus “whitening” the data. This was done by smoothing the spectra with a lowess filter (2 Hz), and then using the FOOOF toolbox (Donoghue et al., 2020) to find the aperiodic signal with data from 2 to 40 Hz, and then subtracting it from the entire spectrum. Finally, power in the theta and alpha band were integrated for both the intact spectra and the burstless spectra, and these values were statistically compared with paired t-tests.

To quantify whether a burst was more likely to occur at a given timepoint for a given trial outcome (Figure 5, Figure 4), we calculated globality (proportion of channels with a burst) for theta and alpha, for each trial outcome at every timepoint. Data was epoched around either the stimulus or response trigger sent from the task computer to the EEG. Timepoints containing artifacts were removed, as were timepoints corresponding to eye closures for Suppl. Figures 4-1 to 7-1. Trials were excluded if more than 50% of the data was missing, for either reason. For the supplementary figures, trials were also excluded if eyes were closed at least 50% of the time during the stimulus window (because the lapse would have been due to the eye-closure). Separately for every trial outcome at every timepoint, globality was averaged across trials, although if there were fewer than 15 trials, data at those timepoints were interpolated. If there were gaps with too few trials larger than 20% of the trial window, then this trial outcome was excluded for that participant; for this reason, the sample size is always provided in the figure. The values for each participant were additionally smoothed over 0.2 s using a lowess filter for visualization purposes. Globality in time for each trial outcome was then z-scored to the mean and standard deviation of globality across the entire EEG session block. Paired t-tests measured deviation from 0, and therefore the average burst globality.

To determine the likelihood of bursts across channels (Figure 6, Figure 7), the proportion of trials containing a burst was calculated for each channel at every timepoint, and this was then averaged within the four windows (Pre: −2-0 s; Stimulus: 0-0.3 s; Response .3-1 s; Post: 2-4 s). The Pre window was chosen a-priori. The stimulus/response windows were split based on reaction times, such that only 1% of RTs were < 0.3 s and thus in the stimulus window. The Post window was chosen a-posteriori to explore further the rebound observed in Figure 4C. These averages were then compared with paired t-tests to the average proportion of bursts for a given channel in a given session block.

### Eye tracking

Eye tracking was done with Pupil Core “glasses” from Pupil Labs. These were eyeglass frames with two rear-facing infra-red cameras. Pupil Player software estimates pupil diameter from the video and provides a confidence value for that estimate; these confidence values were used to determine when participants had their eyes open or closed, with values <.5 considered eyes closed. Consecutive timepoints with confidence values over .5 that lasted less than 50 ms were still considered eyes-closed, and consecutive timepoints under .5 and less than 50 ms long were considered eyes open. Multiple technical failures and poor data quality resulted in data loss such that 6 participants had only 2 BL eye-tracking recordings, one participant had only 1 SD eye tracking, and one participant had no SD eye-tracking.

### Statistics

Paired t-tests were done for every timepoint (Figure 4, 4-1, Figure 5, 5-1) and every channel (Figure 6, 6-1, Figure 7, 7-1), compared to the general likelihood of a burst or eye-closure for that session block (or 0, for z-scored data). All p-values within each plot were adjusted for false-discovery rate (FDR) using the Benjamini-Hochberg method (Benjamini & Hochberg, 1995). The largest t-values are reported in the text as reference.

Hedge’s g effect sizes were calculated to quantify the “meaningfulness” of the t-test results. Using Cohen’s rule of thumb, Hedge’s g around 0.2 is considered small, 0.5 considered medium, and 0.8 large (Becker, 2000; Cohen, 1988). However, given our small sample size it is important to consider what effect sizes we actually had power for. We therefore conducted post-hoc statistical power analysis (MATLAB function *sampsizepwr*) to identify the sample size required for a given effect size, with α = .05 and 1-β=.8. For the full 18 participants, we had statistical power for effect sizes over 0.68, and with 10 participants we had power for effect sizes over 0.94.

When reporting mean values in the text, instead of also providing standard deviations, we indicate the interquartile range (25% and 75% of participants’ values). This can be more informative when results are close to floor or ceiling (e.g. Figure 2B).

### Data and code availability

Data is available upon request. The analysis scripts are on GitHub: https://github.com/snipeso/Lapse-Causes

The cycle-by-cycle analysis library implemented in MATLAB is also on GitHub: https://github.com/HuberSleepLab/Matcycle

## ACKNOWLEDGEMENTS

This study was conducted as part of the SleepLoop Flagship project of Hochschulmedizin Zürich, with additional funding from the Swiss National Science Foundation (320030_179443), Forschungszentrum für das Kind (FZK) of the University Children’s Hospital Zurich, and Hirnstiftung. Professor Hans-Peter Landolt provided the sleep laboratory, Professor Christian Baumann the EEG equipment, Sarah Nadine Meissner and Marc Bächinger the eye tracking equipment. Noa Rieger aided with data collection. Lastly, a special thank you to all of our participants for taking part in this study.

## SUPPLEMENTARY FIGURES

**Figure 4-1:**
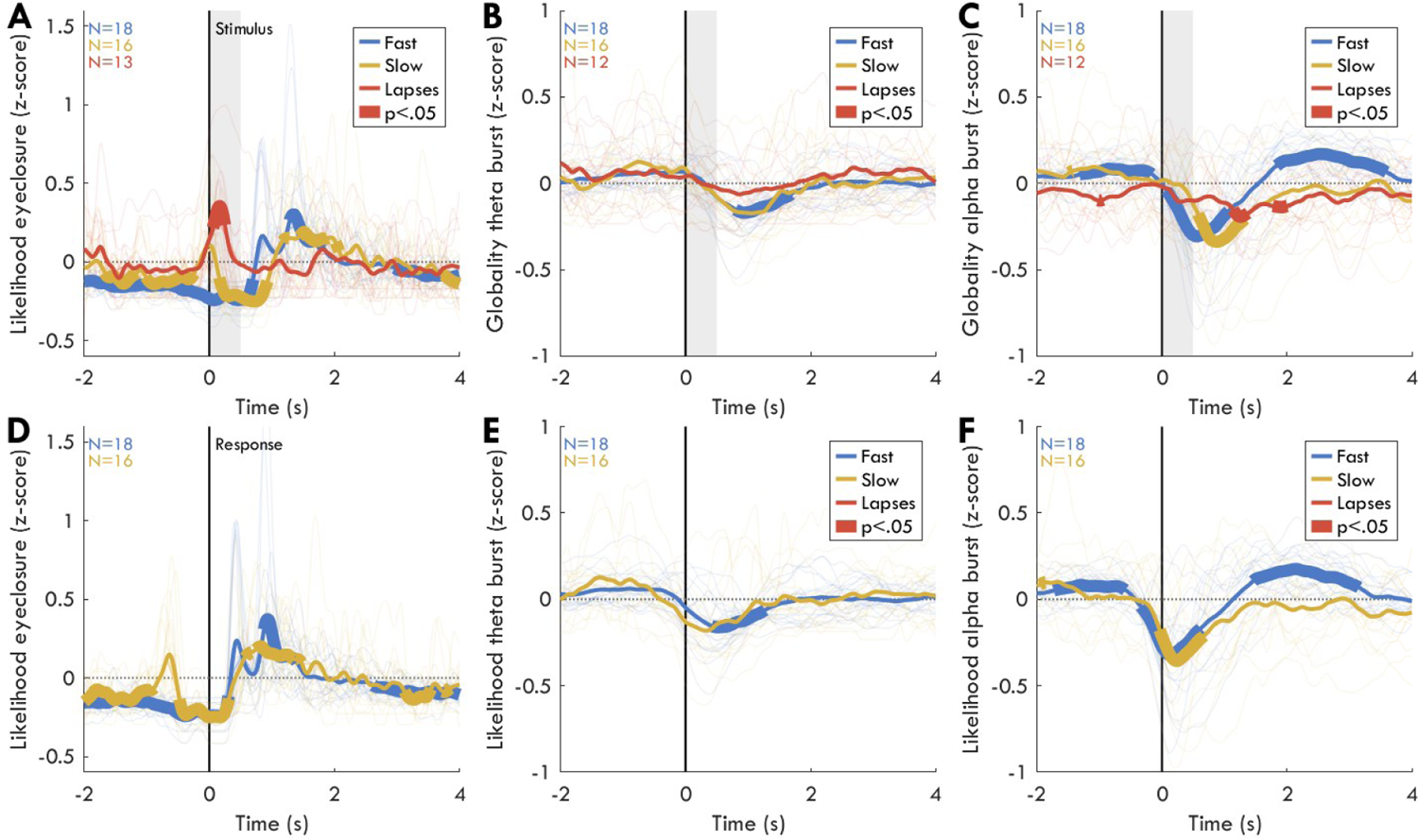
Likelihood of bursts at BL, controlling for eye-closure. Same as Figure 4, excluding trials (B,C,E,F) for which eyes were closed at least 50% during the stimulus window.

**Figure 5-1:**
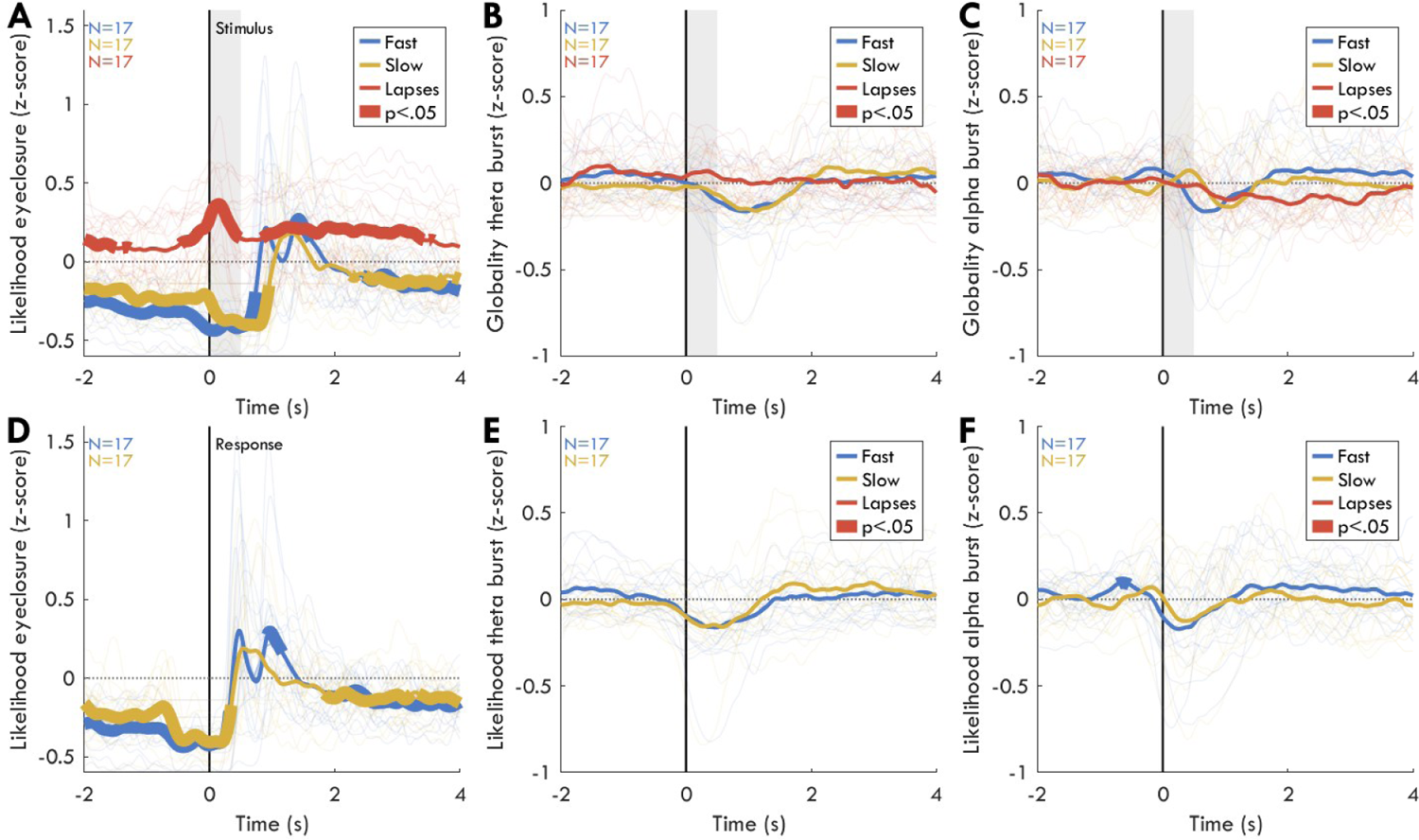
Likelihood of bursts during SD, controlling for eye-closure.

**Figure 6-1:**
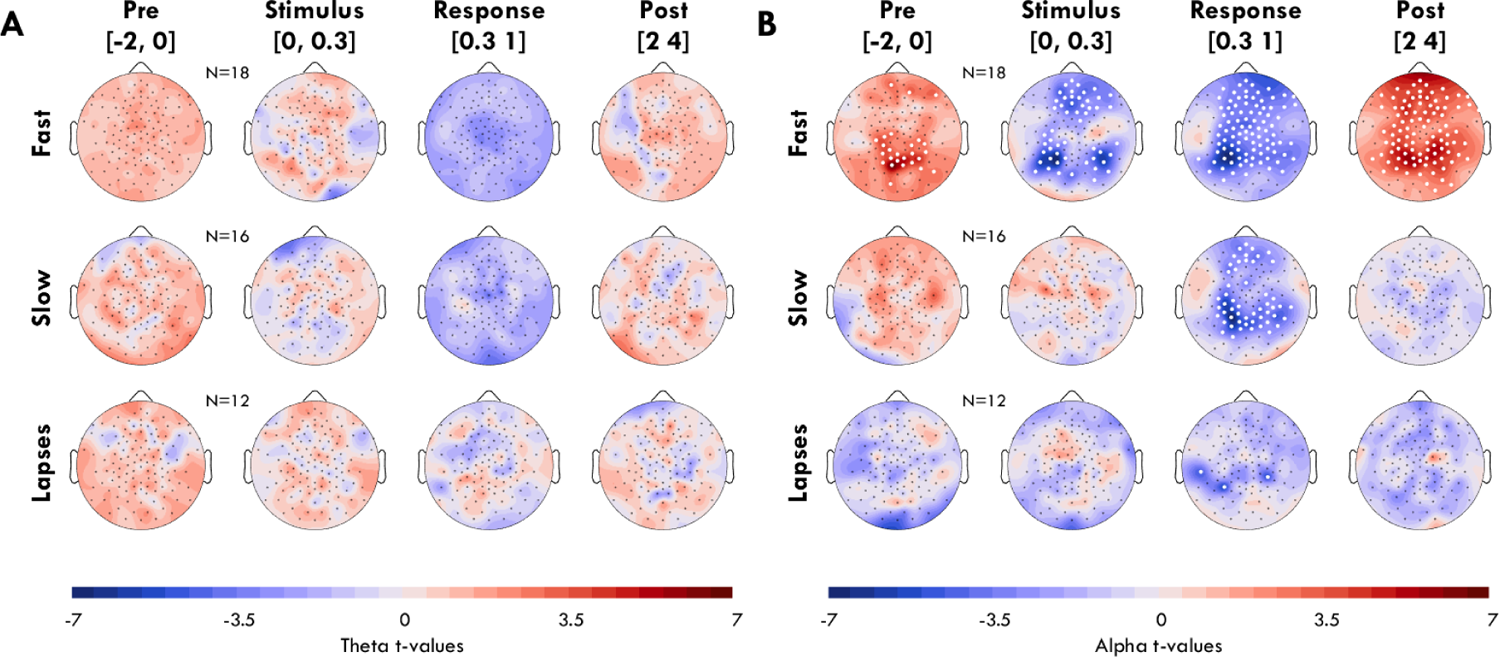
Likelihood of bursts by channel during BL, controlling for eye-closure.

**Figure 7-1:**
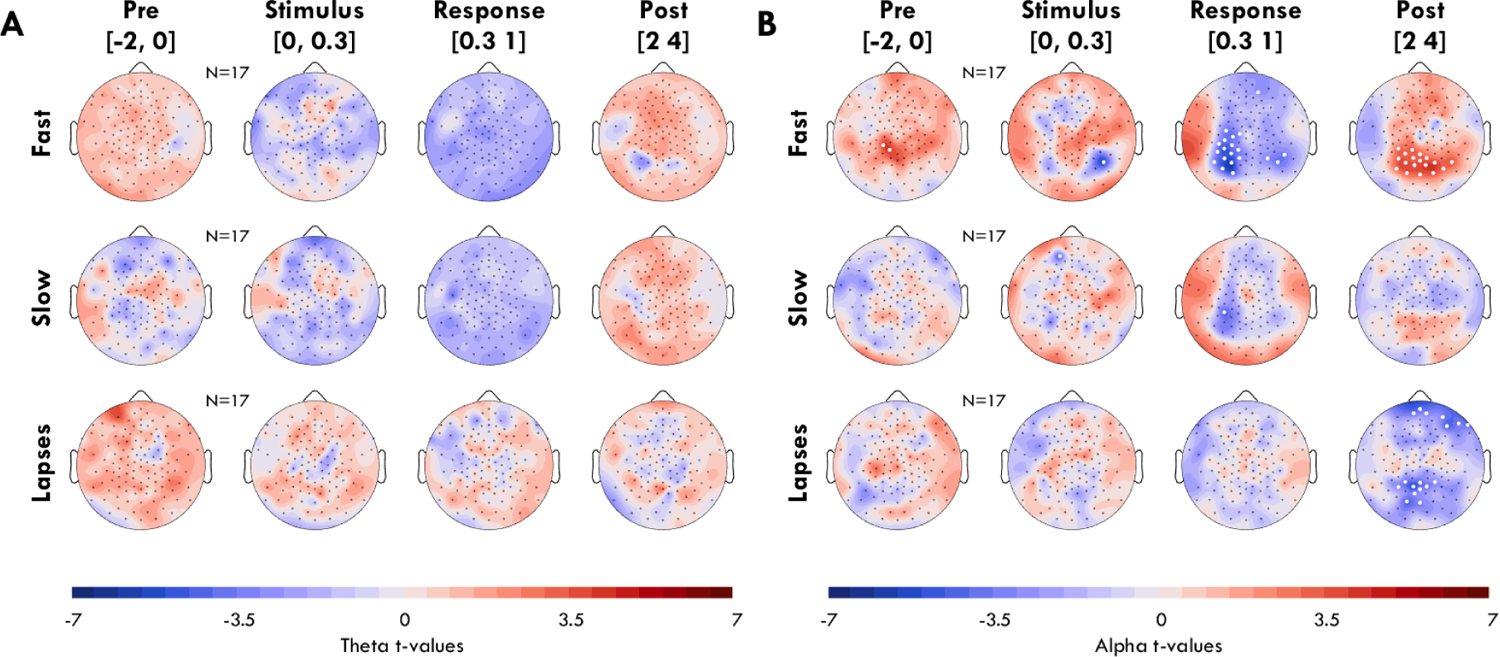
Likelihood of bursts during SD, controlling for eye-closure.

**Figure 8-1:**
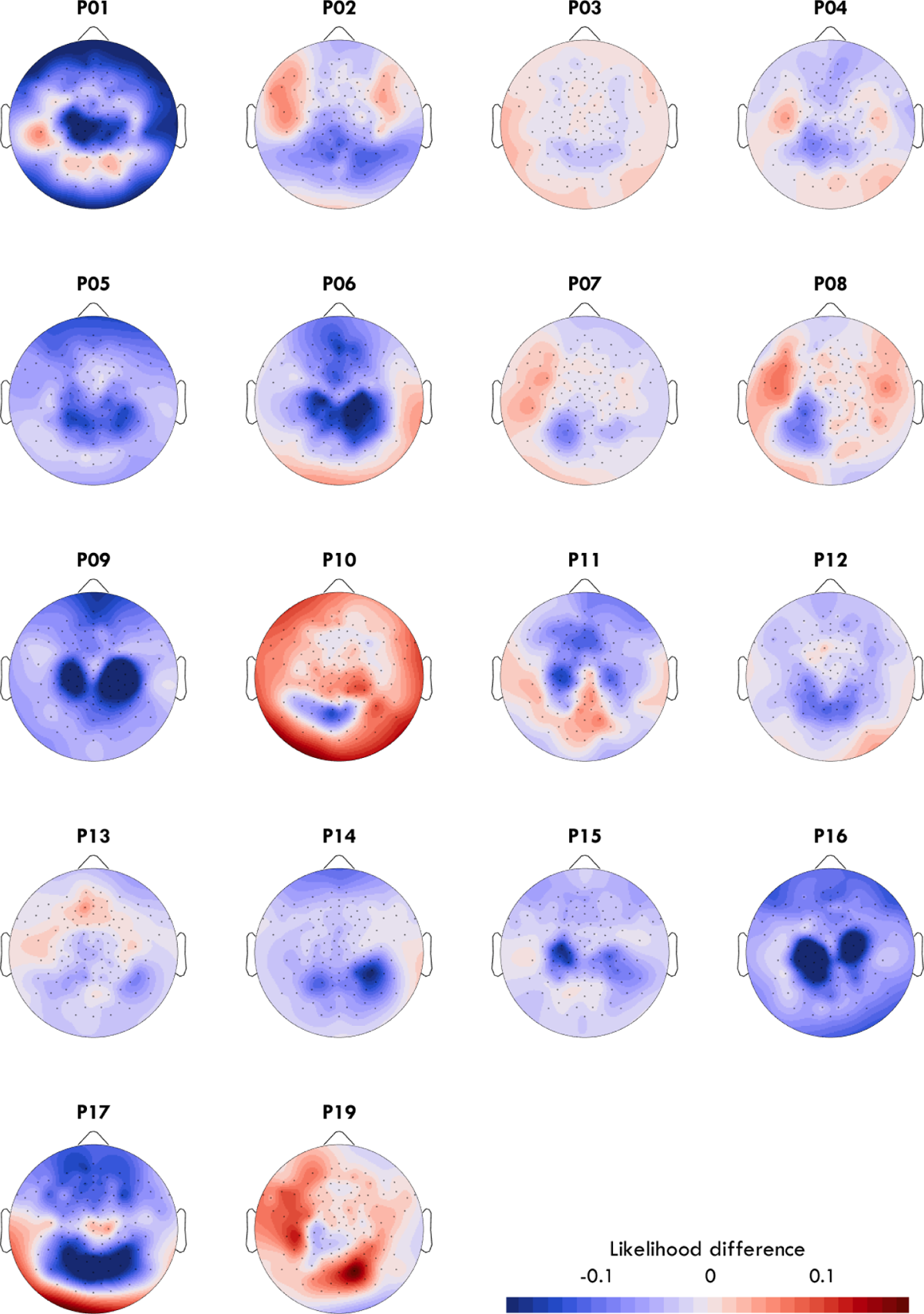
Individual participants change in alpha for fast trials in the response window during BL.

**Figure 8-2:**
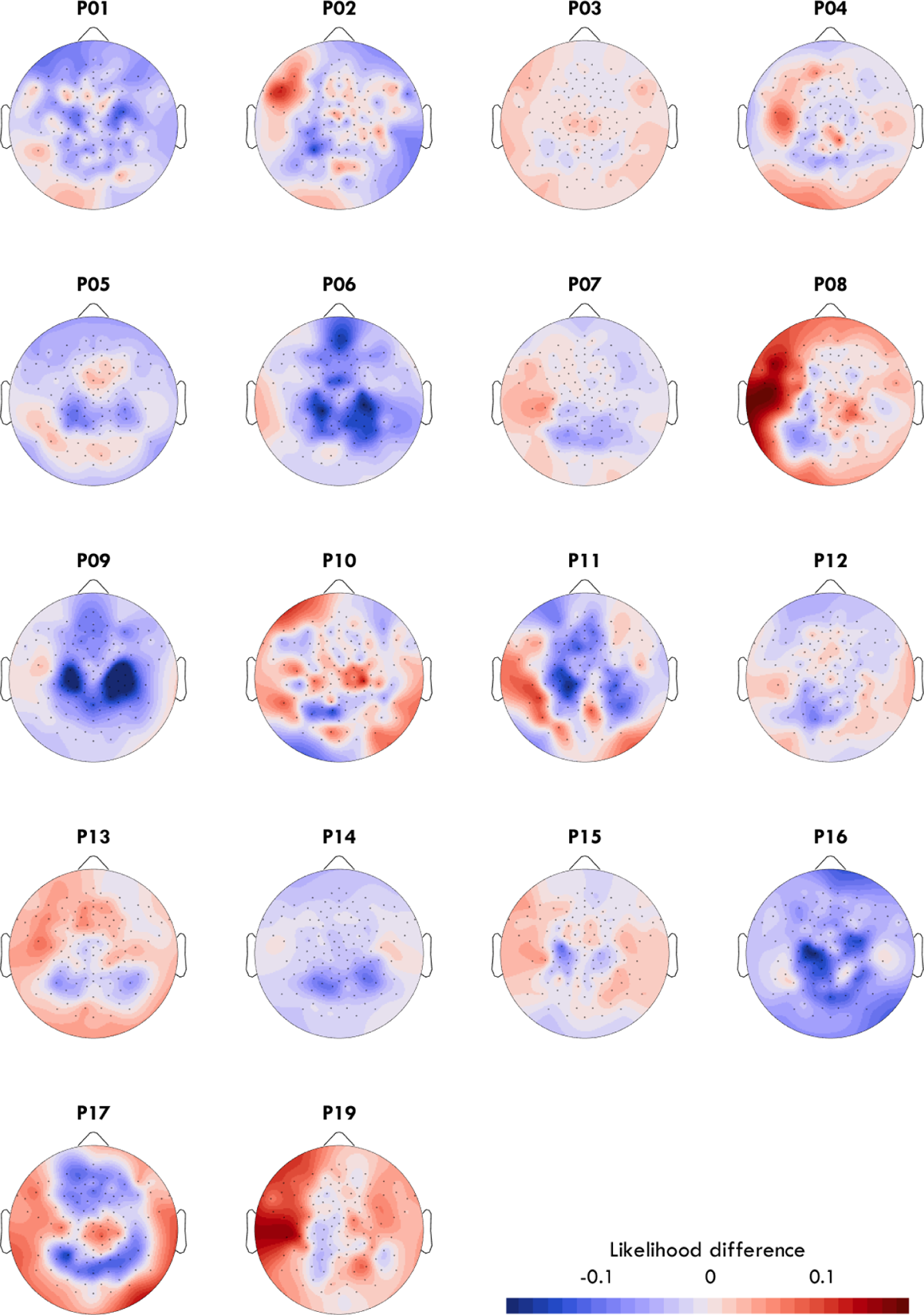
Individual participants change in alpha for fast trials in the response window during SD.

## BIBLIOGRAPHY

Adrian, E., & Matthews, B. (1934). The interpretation of potential waves in the cortex. 81(4).

Aeschbach, D., Matthews, J. R., Postolache, T. T., Jackson, M. A., Giesen, H. A., & Wehr, T. A. (1997). Dynamics of the human EEG during prolonged wakefulness: Evidence for frequency-specific circadian and homeostatic influences. Neuroscience Letters, 239(2–3), 121–124. 10.1016/S0304-3940(97)00904-X

Anderson, C., Wales, A. W. J., & Home, J. A. (2010). PVT Lapses Differ According To Eyes Open, Closed, or Looking Away. Sleep, 33(2), 197–204. 10.1093/sleep/33.2.197

Andrillon, T., Burns, A., Mackay, T., Windt, J., & Tsuchiya, N. (2021). Predicting lapses of attention with sleep-like slow waves. Nature Communications, 12(1), Article 1. 10.1038/s41467-021-23890-7

Basner, M., & Dinges, D. F. (2011). Maximizing sensitivity of the Psychomotor Vigilance Test (PVT) to sleep loss. Sleep, 34(5), 581–591. 10.1093/sleep/34.5.581

Becker, L. A. (2000). Effect size (ES).

Benjamini, Y., & Hochberg, Y. (1995). Controlling the false discovery rate: A practical and powerful approach to multiple testing. Journal of the Royal Statistical Society. Series B (Methodological), 57(1), 289–300. 10.2307/2346101

Bernardi, G., Siclari, F., Yu, I., Zennig, C., Bellesi, M., Ricciardi, E., Cirelli, C., Ghilardi, M. F., Pietrini, P., & Tononi, G. (2015). Neural and behavioral correlates of extended training during sleep deprivation in humans: Evidence for local, task-specific effects. Journal of Neuroscience, 35(11), 4487–4500. 10.1523/JNEUROSCI.4567-14.2015

Cajochen, C., Wyatt, J. K., Czeisler, C. A., & Dijk, D. J. (2002). Separation of circadian and wake duration-dependent modulation of EEG activation during wakefulness. Neuroscience, 114(4), 1047–1060. 10.1016/S0306-4522(02)00209-9

Chee, M. W. L., & Choo, W. C. (2004). Functional Imaging of Working Memory after 24 Hr of Total Sleep Deprivation. Journal of Neuroscience, 24(19), 4560–4567. 10.1523/JNEUROSCI.0007-04.2004

Cohen, J. (1988). *Statistical power analysis for the behavioral sciences* (2nd ed). L. Erlbaum Associates.

Cole, S., & Voytek, B. (2019). Cycle-by-cycle analysis of neural oscillations. Journal of Neurophysiology, 122(2). 10.1152/JN.00273.2019

Dawson, D., & Reid, K. (1997). Fatigue, alcohol and performance impairment. Nature, 388(6639), Article 6639. 10.1038/40775

Delorme, A., & Makeig, S. (2004). EEGLAB: An open source toolbox for analysis of single-trial EEG dynamics including independent component analysis. Journal of Neuroscience Methods, 134(1), 9–21. 10.1016/j.jneumeth.2003.10.009

Dinges, D. F., & Powell, J. W. (1985). Microcomputer analyses of performance on a portable, simple visual RT task during sustained operations. Behavior Research Methods, Instruments, & Computers, 17(6), 652–655. 10.3758/BF03200977

Donoghue, T., Haller, M., Peterson, E. J., Varma, P., Sebastian, P., Gao, R., Noto, T., Lara, A. H., Wallis, J. D., Knight, R. T., Shestyuk, A., & Voytek, B. (2020). Parameterizing neural power spectra into periodic and aperiodic components. Nature Neuroscience, 23(12), Article 12. 10.1038/s41593-020-00744-x

Doran, S. M., Dongen, H. P. A. V., & Dinges, D. F. (2001). Sustained attention performance during sleep deprivation: Evidence of state instability. Archives Italiennes de Biologie, 139(3), Article 3. 10.4449/aib.v139i3.503

Drummond, S. P. A., Meloy, M. J., Yanagi, M. A., Orff, H. J., & Brown, G. G. (2005). Compensatory recruitment after sleep deprivation and the relationship with performance. Psychiatry Research: Neuroimaging, 140(3), 211–223. 10.1016/j.pscychresns.2005.06.007

Fattinger, S., Kurth, S., Ringli, M., Jenni, O. G., & Huber, R. (2017). Theta waves in children’s waking electroencephalogram resemble local aspects of sleep during wakefulness. Scientific Reports, 7(1), Article 1. 10.1038/s41598-017-11577-3

Finelli, L. A., Baumann, H., Borbély, A. A., & Achermann, P. (2000). Dual electroencephalogram markers of human sleep homeostasis: Correlation between theta activity in waking and slow-wave activity in sleep. Neuroscience, 101(3), 523–529. 10.1016/S0306-4522(00)00409-7

Frauscher, B., von Ellenrieder, N., Zelmann, R., Doležalová, I., Minotti, L., Olivier, A., Hall, J., Hoffmann, D., Nguyen, D. K., Kahane, P., Dubeau, F., & Gotman, J. (2018). Atlas of the normal intracranial electroencephalogram: Neurophysiological awake activity in different cortical areas. Brain, 141(4), 1130–1144. 10.1093/brain/awy035

Gibbings, A., Ray, L. B., Gagnon, S., Collin, C. A., Robillard, R., & Fogel, S. M. (2022). The EEG correlates and dangerous behavioral consequences of drowsy driving after a single night of mild sleep deprivation✰. Physiology & Behavior, 252, 113822. 10.1016/j.physbeh.2022.113822

Graw, P., Kräuchi, K., Knoblauch, V., Wirz-Justice, A., & Cajochen, C. (2004). Circadian and wake-dependent modulation of fastest and slowest reaction times during the psychomotor vigilance task. Physiology & Behavior, 80(5), 695–701. 10.1016/j.physbeh.2003.12.004

Hanslmayr, S., Gross, J., Klimesch, W., & Shapiro, K. L. (2011). The role of alpha oscillations in temporal attention. Brain Research Reviews, 67(1), 331–343. 10.1016/j.brainresrev.2011.04.002

Hertig-Godeschalk, A., Skorucak, J., Malafeev, A., Achermann, P., Mathis, J., & Schreier, D. R. (2020). Microsleep episodes in the borderland between wakefulness and sleep. Sleep, 43(1). 10.1093/sleep/zsz163

Horne, J. A., & Reyner, L. A. (1995). Sleep related vehicle accidents. BMJ, 310(6979), 565–567. 10.1136/bmj.310.6979.565

Huang, R.-S., Jung, T.-P., & Makeig, S. (2007). Event-Related Brain Dynamics in Continuous Sustained-Attention Tasks. In D. D. Schmorrow & L. M. Reeves (Eds.), Foundations of Augmented Cognition (pp. 65–74).

Springer. 10.1007/978-3-540-73216-7_8

Hudson, A. N., Van Dongen, H. P. A., & Honn, K. A. (2020). Sleep deprivation, vigilant attention, and brain function: A review. Neuropsychopharmacology, 45(1), Article 1. 10.1038/s41386-019-0432-6

Hung, C. S., Sarasso, S., Ferrarelli, F., Riedner, B., Ghilardi, M. F., Cirelli, C., & Tononi, G. (2013). Local experience-dependent changes in the wake EEG after prolonged wakefulness. Sleep, 36(1), 59–72. 10.5665/sleep.2302

Kirschfeld, K. (2005). The physical basis of alpha waves in the electroencephalogram and the origin of the “berger effect.” Biological Cybernetics, 92(3), 177–185. 10.1007/s00422-005-0547-1

Klimesch, W. (1999). EEG alpha and theta oscillations reflect cognitive and memory performance: A review and analysis. Brain Research. Brain Research Reviews, 29(2–3), 169–195. 10.1016/s0165-0173(98)00056-3

Klimesch, W. (2012). Alpha-band oscillations, attention, and controlled access to stored information. Trends in Cognitive Sciences, 16(12), 606–617. 10.1016/j.tics.2012.10.007

Klimesch, W., Doppelmayr, M., Russegger, H., Pachinger, T., & Schwaiger, J. (1998). Induced alpha band power changes in the human EEG and attention. Neuroscience Letters, 244(2), 73–76. 10.1016/S0304-3940(98)00122-0

Lo, J. C., Groeger, J. A., Santhi, N., Arbon, E. L., Lazar, A. S., Hasan, S., von Schantz, M., Archer, S. N., & Dijk, D. J. (2012). Effects of Partial and Acute Total Sleep Deprivation on Performance across Cognitive Domains, Individuals and Circadian Phase. PLoS ONE, 7(9). 10.1371/journal.pone.0045987

Makeig, S., & Jung, T.-P. (1996). Tonic, phasic, and transient EEG correlates of auditory awareness in drowsiness. Cognitive Brain Research, 4(1), 15–25. 10.1016/0926-6410(95)00042-9

Michels, L., Bucher, K., Lüchinger, R., Klaver, P., Martin, E., Jeanmonod, D., & Brandeis, D. (2010). Simultaneous EEG-fMRI during a working memory task: Modulations in low and high frequency bands. PLoS ONE, 5(4). 10.1371/journal.pone.0010298

Nir, Y., Andrillon, T., Marmelshtein, A., Suthana, N., Cirelli, C., Tononi, G., & Fried, I. (2017). Selective neuronal lapses precede human cognitive lapses following sleep deprivation. Nature Medicine, 23(12), 1474–1480. 10.1038/nm.4433

Ong, J. L., Asplund, C. L., Chia, T. T. Y., & Chee, M. W. L. (2013). Now You Hear Me, Now You Don’t: Eyelid Closures as an Indicator of Auditory Task Disengagement. Sleep, 36(12), 1867–1874. 10.5665/sleep.3218

Pavlov, P. I. (1927). Conditioned reflexes: An investigation of the physiological activity of the cerebral cortex. Annals of Neurosciences, 17(3), 136–141. 10.5214/ans.0972-7531.1017309

Pfeffer, T., Keitel, C., Kluger, D. S., Keitel, A., Russmann, A., Thut, G., Donner, T. H., & Gross, J. (2022). Coupling of pupil- and neuronal population dynamics reveals diverse influences of arousal on cortical processing. eLife, 11, e71890. 10.7554/eLife.71890

Rihs, T. A., Michel, C. M., & Thut, G. (2007). Mechanisms of selective inhibition in visual spatial attention are indexed by α-band EEG synchronization. European Journal of Neuroscience, 25(2), 603–610. 10.1111/j.1460-9568.2007.05278.x

Rihs, T. A., Michel, C. M., & Thut, G. (2009). A bias for posterior α-band power suppression versus enhancement during shifting versus maintenance of spatial attention. NeuroImage, 44(1), 190–199. 10.1016/j.neuroimage.2008.08.022

Sauseng, P., Klimesch, W., Stadler, W., Schabus, M., Doppelmayr, M., Hanslmayr, S., Gruber, W. R., & Birbaumer, N. (2005). A shift of visual spatial attention is selectively associated with human EEG alpha activity. European Journal of Neuroscience, 22(11), 2917–2926. 10.1111/j.1460-9568.2005.04482.x

Scheeringa, R., Bastiaansen, M. C. M., Petersson, K. M., Oostenveld, R., Norris, D. G., & Hagoort, P. (2008). Frontal theta EEG activity correlates negatively with the default mode network in resting state. International Journal of Psychophysiology, 67(3), 242–251. 10.1016/j.ijpsycho.2007.05.017

Scheeringa, R., Petersson, K. M., Kleinschmidt, A., Jensen, O., & Bastiaansen, M. C. M. (2012). EEG Alpha Power Modulation of fMRI Resting-State Connectivity. Brain Connectivity, 2(5), 254–264. 10.1089/brain.2012.0088

Scheeringa, R., Petersson, K. M., Oostenveld, R., Norris, D. G., Hagoort, P., & Bastiaansen, M. C. M. (2009). Trial-by-trial coupling between EEG and BOLD identifies networks related to alpha and theta EEG power increases during working memory maintenance. NeuroImage, 44(3), 1224–1238. 10.1016/j.neuroimage.2008.08.041

Schomer, D. L., & Silva, F. H. L. da. (2011). Niedermeyer’s Electroencephalography: Basic Principles, Clinical Applications, and Related Fields. Lippincott Williams & Wilkins.

Smith, J. R. (1938). The Electroencephalogram During Normal Infancy and Childhood: I. Rhythmic Activities Present in the Neonate and Their Subsequent Development. The Pedagogical Seminary and Journal of Genetic Psychology, 53(2), 431–453. 10.1080/08856559.1938.10533820

Snipes, S., Krugliakova, E., Meier, E., & Huber, R. (2022). The theta paradox: 4-8 Hz EEG oscillations reflect both sleep pressure and cognitive control. Journal of Neuroscience. 10.1523/JNEUROSCI.1063-22.2022

Snipes, S., Meier, E., Meissner, S. N., Landolt, H.-P., & Huber, R. (2023). How and when EEG reflects changes in neuronal connectivity due to time awake. iScience, 26(7). 10.1016/j.isci.2023.107138

Strijkstra, A. M., Beersma, D. G. M., Drayer, B., Halbesma, N., & Daan, S. (2003). Subjective sleepiness correlates negatively with global alpha (8-12 Hz) and positively with central frontal theta (4-8 Hz) frequencies in the human resting awake electroencephalogram. Neuroscience Letters, 340(1), 17–20. 10.1016/S0304-3940(03)00033-8

Van Dongen, H. P. A., Maislin, G., Mullington, J. M., & Dinges, D. F. (2003). The Cumulative Cost of Additional Wakefulness: Dose-Response Effects on Neurobehavioral Functions and Sleep Physiology From Chronic Sleep Restriction and Total Sleep Deprivation. Sleep, 26(2), 117–126. 10.1093/sleep/26.2.117

Vyazovskiy, V. V., Olcese, U., Hanlon, E. C., Nir, Y., Cirelli, C., & Tononi, G. (2011). Local sleep in awake rats. Nature, 472(7344), 443–447. 10.1038/nature10009

Vyazovskiy, V. V., & Tobler, I. (2005). Theta activity in the waking EEG is a marker of sleep propensity in the rat. Brain Research, 1050(1–2), 64–71. 10.1016/j.brainres.2005.05.022

